# Photoswitchable optoGPCRs for reversible control of Gs signalling and arrestin recruitment

**DOI:** 10.64898/2026.07.01.735182

**Authors:** Deborah Walter, Richard McDowell, Polina Isaikina, Andreea Pantiru, Hiruttika Ravimohan, Xavier Deupi, Robert J. Lucas, Gebhard F. X. Schertler

## Abstract

OptoGPCRs are light-activatable G protein-coupled receptors (GPCRs) used to optogenetically control physiological processes. Most existing optoGPCRs are based on monostable opsins, which are limited by photobleaching and irreversibility. The bistable jumping spider rhodopsin 1 (JSR1) carrying the single point mutant S199F introduces a ∼150 nm spectral separation between the active and inactive states, enabling bidirectional control with distinct wavelengths of light. Here, we show that JSR1-S199F robustly and reversibly recruits arrestin and modulates Gq/i protein activity. We then engineered JSR1-S199F-based optoGPCRs with Gs protein activity, guided by AlphaFold-predicted receptor-miniGs complexes, to expand the limited repertoire of bistable Gs-coupled optoGPCRs. Specifically, we present optoDRD1, a chimeric optoGPCR that redirects the native Gq/i protein activity of JSR1 towards the Gs pathway of the dopamine D1 receptor (DRD1). Through systematic screening of intracellular domain combinations, we identified a chimeric configuration with the highest absolute cAMP output comprising ICL2, ICL3, helix 8, and the C-terminus from DRD1. The resulting optoGPCR is activated by violet light (λ_max_ = 397 nm) and deactivated by green light (λ_max_ = 531 nm) at physiologically relevant light intensities. A single violet light pulse drives sustained Gs signalling for several hours, while green light rapidly deactivates the receptor at any chosen time point, enabling repeatable switching between active and inactive states over multiple cycles. OptoDRD1 closely mimics wild-type DRD1 in both cAMP signalling kinetics and G protein selectivity, with a signal amplitude comparable to native Gs-coupled receptors, including JellyOp. We further demonstrate optoDRD1’s utility for optogenetic control of Gs-regulated processes *in vitro*, including insulin secretion in human β-cells and signalling modulation in a neuronal cell line. The biochemical stability and known structure of JSR1 make it a robust scaffold for this rational engineering and for future biophysical characterization. Together, optoDRD1 and JSR1-S199F expand the optoGPCR toolkit and open new opportunities for dissecting dopaminergic signalling, Gs-mediated physiology, and GPCR signalling pharmacology.

## Introduction

Optogenetics is a technology that enables precise spatiotemporal control of cellular activity through a combination of genetic and optical methods. The field largely relies on the use of microbial light-sensitive ion channels, known as Type I opsins, e.g., algal channelrhodopsin, to control membrane potential and modulate neuronal activity^1,2^. In contrast, animal opsins, also known as Type II opsins, belong to the large and diverse family of G protein-coupled receptors (GPCRs). GPCRs are a cornerstone of biomedical research due to their pivotal roles as mediators of cellular signalling. They typically respond to extracellular stimuli, such as neurotransmitters and hormones, and represent one of the largest drug target families^3^. Activation of GPCRs typically results in the activation of heterotrimeric G proteins. These can be classified into four families, according to the nature of the α subunit: Gs, Gi/o/t, Gq/11 and G12/13. Each one of them can alter the activity of effector enzymes directly or through second messengers such as cAMP or calcium. Thus, animal opsins are light-sensitive GPCRs able to modulate diverse cellular signalling cascades that regulate gene expression and cell fate, which makes them promising candidates for developing the next generation of optogenetics tools.

Opsins bind the 11-*cis* isomer of the retinal chromophore covalently via a protonated Schiff base linkage to lysine K^7.43^ (superscripts indicate the GPCRdb residue numbering scheme^4^). Upon absorption of a photon, 11-*cis*-retinal isomerises to the all-*trans* isomer. In monostable opsins, such as visual rhodopsins, isomerisation leads to hydrolysis of the Schiff base and subsequent retinal release. Thus, monostable opsins must be reconstituted with 11-*cis*-retinal to recover photosensitivity, limiting their use for *in vivo* optogenetic applications. In contrast, bistable opsins, mostly invertebrate opsins and a few non-visual vertebrate opsins, do not release the isomerised all-*trans*-retinal. Instead, they can absorb another photon to isomerise back to 11-*cis*-retinal. Therefore, bistable opsins are natural molecular photoswitches, alternating between inactive and active states by absorbing light at defined wavelengths. Bistable opsins from various organisms have been used to develop optogenetic tools^5–11^. The bistable jumping spider rhodopsin 1 (JSR1) from *Hasarius adansoni* shows particular promise as it is exceptionally stable, expresses well and easily regenerates the retinal as shown in biochemical studies^12,13^. In the wild-type receptor the absorbance spectra of the inactive state (11-*cis*-retinal) and the active photoproduct (all-*trans*-retinal) overlap (λ_max_ = 535 nm). However, the S199F (S^45.49^F) mutation on extracellular loop two perturbs the hydrogen-bonding network stabilizing the inactive state, resulting in two spectrally well-separated states, where the inactive state has λ_max_ ≈ 385 nm and the active state has λ_max_ ≈ 540 nm^14^. This spectral separation allows JSR1-S199F to be robustly and reversibly switched between states with violet and green light, enabling precise optical control of receptor activity^15^.

Many optogenetic applications necessitate adjusting intrinsic opsin properties, such as colour tuning or G protein signalling selectivity. These alterations yield engineered light-sensitive GPCRs (optoGPCRs) with tailored signalling profiles^6,15,16^. The most widely used approach for altering G protein selectivity is the generation of chimeric optoGPCRs. These chimeras combine an animal opsin as the light-sensitive core, with intracellular domains grafted from another GPCR, as the intracellular surface is crucial for selective G protein recognition. Pioneering work was conducted with monostable bovine rhodopsin and the human β_2_ adrenergic receptor (β2AR)^16^. Later strategies incorporated available structural information, improving the design of optoGPCRs^17^. The first bistable chimeric optoGPCR combined human melanopsin with the glutamate receptor mGluR6^18^.

Most of the currently available optoGPCR tools activate Gi or Gq proteins. JSR1 is no exception and signals via both human Gi and Gq^19^. In contrast, Gs-coupled opsins are rare^20–22^, and engineered bistable optoGPCR tools remain limited. A naturally bistable Gs-coupled opsin was recently reported in a cnidarian species^23^. However, tools based on native mammalian Gs-coupled GPCR pharmacology with reversible temporal control have not yet been described. Since Gs pathways regulate diverse physiological functions, such as motor activity, cognition and learning^24^, cardiovascular control^25^, and blood glucose regulation^26^, expanding the repertoire of Gs-coupled optoGPCRs represents an important step in diversifying the functional applications of optogenetic tools.

Here, we report cellular signalling data, G protein selectivity, and arrestin recruitment properties of JSR1 and its S199F mutant. Furthermore, we describe chimera design with the help of AlphaFold (AF) and optoDRD1, a chimeric optoGPCR generated by combining the light-sensitive core of JSR1-S199F with the intracellular domains of the human dopamine D1 receptor (DRD1), a Gs-coupled receptor abundantly expressed in striatal neurons^24^. We show that optoDRD1 exhibits a cAMP signalling time-course comparable to that of wild-type DRD1, with light-dependent activation (385 nm) and deactivation (525 nm) of Gs signalling in HEK293T and Neuro-2a cells. Thus, optoDRD1 provides a genetically encoded, light-controlled tool for precise spatiotemporal control of Gs signalling, complementing existing chemogenetic approaches such as Gs-coupled designer receptors exclusively activated by designer drugs (DREADDs)^27^.

## Results

### Comparison of signalling properties of JSR1 wild-type (WT) and the JSR1-S199F mutant

JSR1 is the bistable rhodopsin that serves as the engineering scaffold for our optoGPCR designs. JSR1 natively couples to a jumping spider Gq protein^28^, and it has previously been shown to activate both the human Gi and Gq pathway *in cellulo*^15,29^ and in purified protein samples^19^. To confirm that recombinantly expressed JSR1-S199F signals similarly to JSR1-WT through both human Gi and Gq *in cellulo*, we first assessed Gq-mediated intracellular Ca^2+^ elevation using the bioluminescent indicator aequorin. Both JSR1-WT and JSR1-S199F elicit comparable Ca^2+^ responses of similar amplitude **(Fig. S1a-b)** as described previously^14^. We additionally assessed G protein selectivity between Gi, Gq and Gs using the bioluminescence resonance energy transfer (BRET) G protein activation assay (**Fig. 1a**)^30,31^. Neither JSR1-WT nor JSR1-S199F elicited a detectable Gs response. JSR1-S199F showed robust activation of both Gi and Gq proteins (**Fig. 1a-b**). In contrast, JSR1-WT displayed robust Gi activation (Δratio = 0.2) but a markedly reduced Gq response (Δratio = 0.05), compared to JSR1-S199F (Δratio = 0.4, **Fig. 1a**). Nevertheless, the association kinetics showed Gq binding was approximately twice as fast as Gi for both variants **(Fig. S1c-d)**, suggesting that the S199F mutation enhances the amplitude of Gq signalling without markedly affecting binding kinetics^31^. These results show that even a single point mutation (S199F) in the ligand binding pocket can allosterically modulate the G protein binding profile. The BRET G protein activation assay further confirmed that the JSR1-S199F activity can be switched off with green light and subsequently reactivated with violet light *in cellulo* (**Fig. 1b**) as shown previously *in vivo*^15^. Thus, JSR1-S199F can be repeatedly and reversibly activated (385 nm) and deactivated (525 nm) with distinct wavelengths of light. We further confirmed that purified JSR1-S199F, reconstituted with 9-*cis*-retinal, exhibits a distinct spectral separation of the ground state (λ_max_ ≈ 365 nm) and the active photoproduct (λ_max_ ≈ 540 nm, **Fig. 1c**). Subsequent illumination with green light shifts the absorbance maximum back to the ground state, consistent with the previously described photochemical properties^12,14,32^. **Fig. 1d** illustrates the strategy of our design, using the light-switching mechanism of the JSR1-S199F colour-tuning single point mutation^14^ and altering its signalling properties from Gq/Gi to Gs by grafting intracellular domains of human Gs coupled receptors.

**Figure 1.**
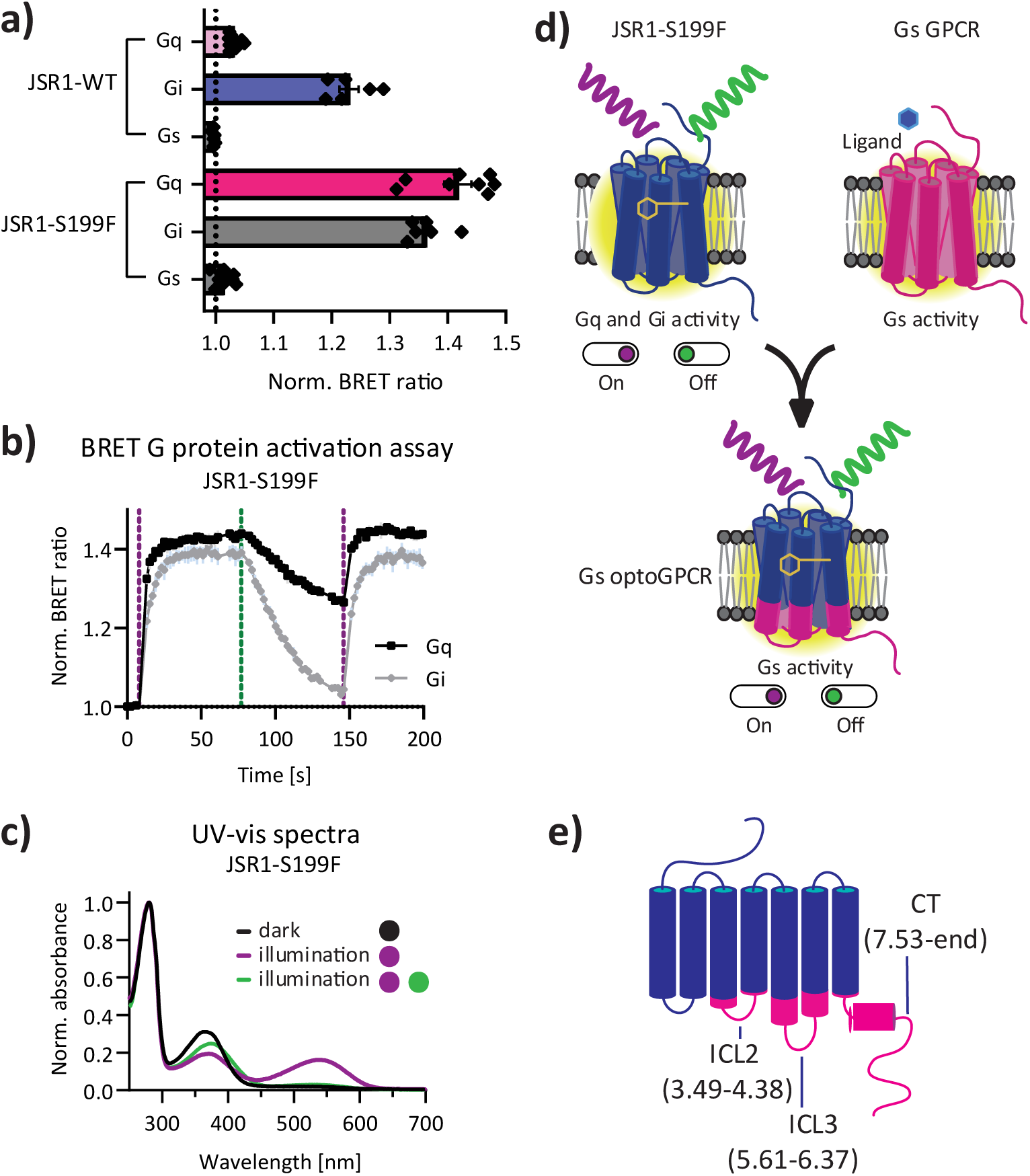
Signalling and photochemical properties of the JSR1-WT and JSR1-S199F mutant before chimera generation. **a)** Normalised BRET ratio for Gq, Gi and Gs activation by JSR1-WT and JSR1-S199F in HEK293T cells (mean± SEM). n = 3 biological replicates (3 technical replicates each) for JSR1-WT Gq and JSR1-S199F Gq and Gs; n = 2 biological replicates (3 technical replicates each) for JSR1-WT Gi and Gs and JSR1-S199F Gi. **b)** Time-resolved signalling profiles of JSR1-S199F in a BRET G protein activation assay in HEK293T cells, showing Gq and Gi activity after illumination for 1 s at dashed lines with 385 nm (10^15.5^ photons cm^-2^), 525 nm (10^15.5^ photons cm^-2^), and 385 nm (10^15.5^ photons cm^-2^). The data represent mean ± SEM of two biological replicates (n=6). Comparison to JSR1-WT and kinetic analysis are shown in Fig. S1. **c)** UV-vis spectra of purified JSR1-S199F reconstituted with 9-*cis*-retinal. The dark state shows λ_max_ ≈ 365 nm (black line). Illumination with violet light (365 nm, ∼10^17^ photons cm^-2^) partially shifts the chromophore absorbance to λ_max_ ≈ 540 nm (purple) while retaining some absorbance at 365 nm, and subsequent illumination with green light (525 nm, ∼10^17^ photons cm^-2^) shifts the absorbance back to the inactive state absorbance of λ_max_ ≈ 385 nm (green). n = 1, spectra from n = 3 independent purifications are in Fig. S1f. **d)** Depiction of the engineering strategy. The goal is to create a Gs signalling optoGPCR with bidirectional control of activity. The data for chimeric optoGPCRs will be shown in later figures. **e)** Scheme of the JSR1 chimeras in which ICL2, ICL3, and the C-terminus (CT) are exchanged with the sequence of a Gs-receptor (pink) to redirect its native signalling properties towards Gs signalling.

### AlphaFold- and structure-guided design of optoGPCRs

Several engineered light-sensitive versions of diffusible ligand-binding GPCRs – optoGPCRs or optoXRs – have been reported^9,16,17,33,34^. However, extending these designs to new receptor combinations is non-trivial and often requires extensive screening and optimization. Our engineering strategy builds on these previous approaches and incorporates experimental and predicted structures to assess the quality of the designed chimeras.

Our initial designs were based on JSR1-WT, as the experimental structures of both, the inactive and G protein-bound active states are available^12,19^. We designed a series of chimeras by replacing the G protein binding interface of JSR1 with that from the Gs-coupled β2-adrenergic (β2AR), dopamine D1 (DRD1), adenosine A2B (AA2BR), serotonin 7 (5HTR7), trace amine-associated 9 (TAAR9), luteinizing hormone (LSHR), and JellyOp receptors. The structures of all these receptors (except JellyOp) have been solved in complex with Gs protein (PDB IDs: 8GG0, 7JV5, 8HDP, 7XTC, 8IW9, 7FIH). JellyOp was included as the best-characterised native Gs-coupled opsin and since its intracellular loop 3 (ICL3) has previously been used to generate Gs-coupled chimeras with other bistable opsins^9,35^. Therefore, we included one such chimera of JSR1 with the ICL3 of JellyOp (5.60-6.34) as a control^35^. For the other designs, we selected a diverse set of receptors with longer ICLs and C-termini (DRD1, β2AR and 5HT7R), as well as receptors with short sequences (AA2BR, TAAR9 and LSHR, **Table S1**).

Using the GPCR-G protein interface maps from the GPCRdb^36,37^, we extracted the contacts between the receptor and G protein for all available experimental structures of human Class A Gs-coupled receptors, excluding fusion constructs (n = 186). The contacts formed by different Gs-coupled receptors with the Gs protein are heterogeneous. Considering only interactions present in at least 20% of the structures, we identified one conserved contact in intracellular loop 1 (ICL1), ten in ICL2, 14 in ICL3 and four at the C-terminus (CT) at the junction of transmembrane helix 7 and helix 8 **(Fig. S2)**. Possible contacts established by the flexible receptor CT are typically not resolved in the structures and therefore not present in this analysis. Nonetheless, the C-terminus has been shown to be involved in receptor signalling^38^, including receptor autoinhibition^39^, GPCR kinase (GRK) and arrestin recruitment^40^, and receptor localization^41^. As a result of this analysis, for our optoGPCR chimera design we decided to replace ICL2, ICL3 and the CT and include as many identified contact sites as possible (**Fig. 1e**). For simplicity we call the exchanged regions ICL2, ICL3 and CT although they also contain part of the ordered alpha-helical domains, as apparent from their GPCRdb number 3.49 (helix 3 position 49), 34.55 (loop between helix 3 and 4). This strategy is similar to previous approaches used for other chimeric opsins^16,17,34^ and offers a more complete recreation of G protein signalling than exchanging ICL3 alone, which has previously been shown to elicit some Gs signalling response^35^. For the JSR1-β2AR chimera, we used the optimised design described in Tichy et al.^17^ between β2AR and bovine rhodopsin. Although ICL1 harbours one conserved G protein contact, it was excluded from the design as previous studies have reported that its incorporation into the chimera can destabilise the receptor and reduce expression levels^16,42^. The sequences of all receptors used for chimera designs were aligned, and the loop regions were exchanged between ICL2 (3.49-4.38), ICL3 (5.60/61-6.37/40), and the CT (7.53-end) including ICL4 and helix 8 (**Fig. 1e, Table S2)**. The precise grafting positions and a sequence alignment of the JSR1-DRD1 chimera are shown **in Fig. S2d**. To preserve structural integrity, we ensured overlap at the domain junctions when possible. Therefore, we swapped the amino acid sequences directly after conserved motifs involved in G protein binding^43^ and present in JSR1 and the Gs receptors.

Using AF, we generated models of the active state of the chimeric receptors by predicting their complexes with miniGs, an engineered minimised form of Gs^44^, testing several slightly shifted domain junctions per design. Designs were prioritised when the grafting points retained pLDDT > 70 (predicted local distance difference test, 0–100; **Fig. 2a**) and the interface confidence scores were among the highest of the designs tested. We also required that the grafted regions in the AF models resembled the experimental structures. No fixed threshold was applied.

**Figure 2.**
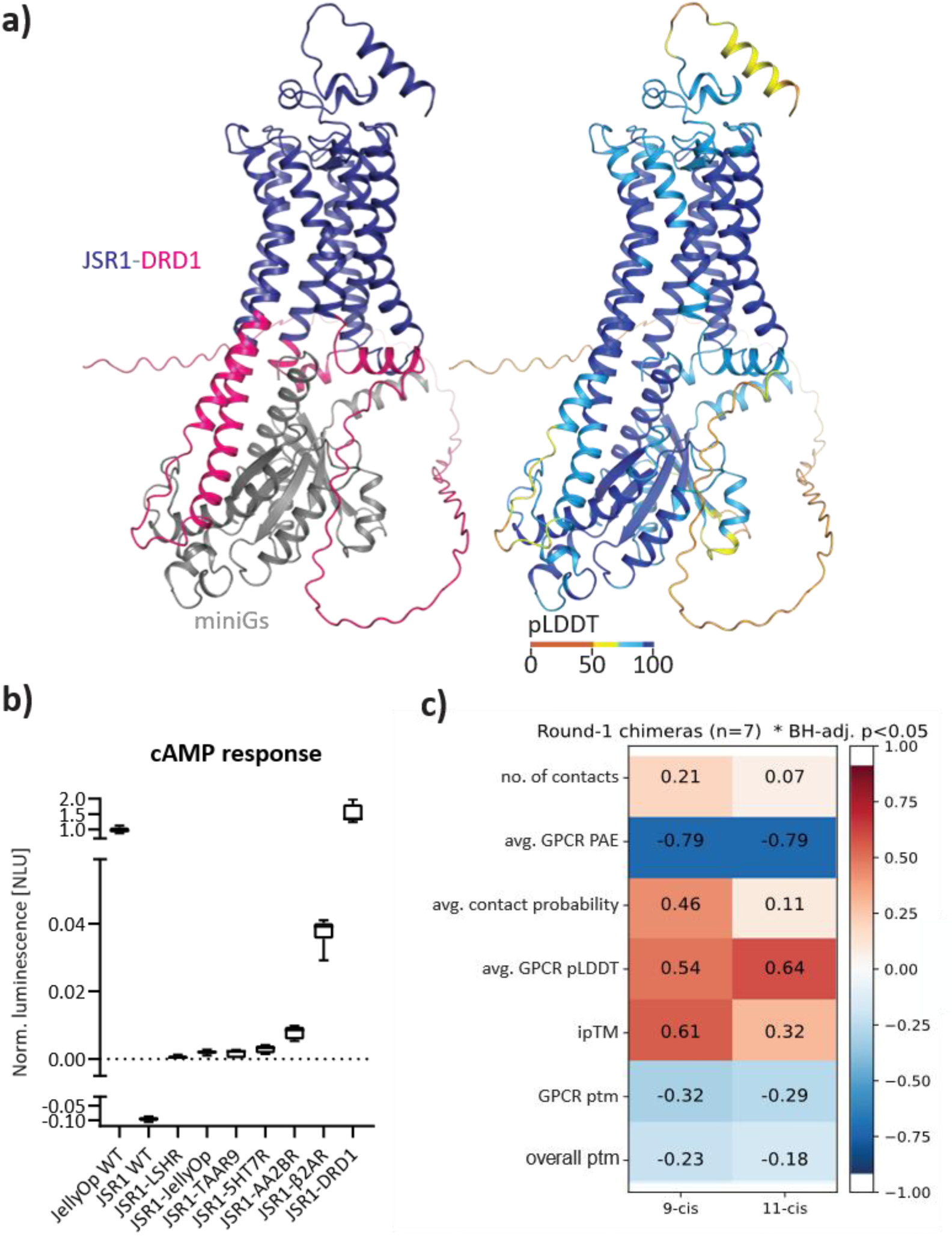
Chimera design strategy and evaluation. **a)** AF model of the JSR1-DRD1-miniGs complex. On the left, the model is colour-coded according to the sequence (dark blue: JSR1, pink: DRD1, grey: miniGs), and on the right, the model is colour-coded according to pLDDT scores (confidence of the AF prediction). **b)** Maximal amplitude of GloSensor cAMP response of the different chimeras expressed in HEK293T cells supplemented with *9-cis*-retinal and stimulated with the OptoWell (519 nm, ∼10^17^ photons cm^-2^). The values are displayed after subtraction of the dark signal and normalization to the JellyOp response (n = 6, from two biological replicates). Chimeras and JSR1-WT were measured with 1 µM forskolin; JellyOp without forskolin. Amplitudes are therefore not directly comparable between these groups. The limits of the boxes represent 25^th^ to 75^th^ percentiles and the whiskers the minimum and maximum values. Data obtained with 11-*cis*-retinal are shown in Fig. S4 and representative time-resolved data are presented in Fig. S5. **c)** Spearman correlation matrix between AF interface-confidence metrics and the maximum cAMP amplitude of cells supplemented with 9-*cis*- or 11-*cis*-retinal (data in Figs. S4–S5). Colour and values indicate Spearman’s ρ; no correlation reached Benjamini-Hochberg (BH) adj. *p* < 0.05. Correlations were calculated across all constructs with data for both conditions (n = 7).

We evaluated each model with four confidence metrics at the GPCR-G protein interface. The interface predicted template modelling score (ipTM, 0–1) reports the confidence in the relative placement of the two chains. The contact probability reports the predicted likelihood that residues from the two chains are in contact (AF defines this by a distance < 8 Å), we considered contacts with a probability > 0.2 as part of the interface. Of the extracted interface residues, we calculated the average predicted aligned error (PAE, in Å), which estimates the positional error between residue pairs. Lower values indicate a more confidently predicted interface. Additionally, the average pLDDT of interface residues reports per-residue local confidence and was used to verify that the interface architecture was intact **(Tables S3–S4)**. These scores estimate model confidence at the binding interface, not binding affinity or signalling efficacy, and are only informative for the relative ranking of related designs.

### Evaluation of signalling properties of optoGPCR designs

Immunocytochemistry assays in HEK293T cells confirmed that all chimeric proteins were expressed **(Fig. S3)**. As these assays were performed on permeabilized cells, they report on total cellular immunoreactivity and localization pattern rather than quantitative plasma-membrane trafficking. Using a GloSensor cAMP assay, we observed that all chimeras lost the native Gi activity of JSR1-WT, as shown by their inability to suppress forskolin-induced cAMP (**Fig. 2b**). Instead, most chimeras, except for JSR1-LSHR, elicited a modest light-induced increase in cAMP, with amplitudes and durations of signal that vary for each optoGPCR **(Fig. S4-5)**. In the screening conditions, JSR1-DRD1 showed the strongest cAMP response, approximately 30% higher than JellyOp (**Fig. 2b**). As forskolin was added for JSR1-WT and for the chimeras but not for JellyOp **(Fig. S5)**, this comparison is not matched, and a controlled head-to-head comparison is shown in **Fig. 5a, 4f and S14**. As comparison all the other chimeras are reaching cAMP levels of below 4% compared to JellyOp’s response in 9-*cis*-retinal (**Fig. 2b**). When supplemented with 11-*cis*-retinal, most chimeras produced a higher response, with the highest responding chimera (JSR1-β2AR) reaching 30% of JellyOp’s response **(Fig. S4)**. This is nevertheless much lower than the JSR1-DRD1 response with 9-*cis*-retinal.

To rationalise the activity of the designed chimeras, we analysed the AF-predicted models, focusing on confidence values at the receptor-G protein interface. In all the models, miniGs inserts deeply into the open intracellular cavity of the transmembrane bundle (**Fig. 2a**), consistent with experimentally resolved structures. Interface confidence scores decreased when a receptor was modelled with a G protein to which it is not natively coupled, as illustrated by JSR1-WT, a Gi/q-coupled receptor, modelled with miniGs **(Fig. S6)**. To focus on the interaction surface, we computed the average contact probability and the predicted alignment error (PAE) of receptor–G protein residue pairs with a contact probability above 0.2 **(Table S3)**. Among all chimeras, the AF prediction for the JSR1-DRD1–Gs complex yielded one of the highest confidence scores.

To determine whether these AF-derived metrics relate to the signalling abilities of the constructs, we calculated Spearman rank correlations between the AF confidence metrics and the maximum cAMP amplitude measured with 9-cis- or 11-cis-retinal supplementation (**Fig. 2c**; underlying data in **Figs. S4–S5, Table S3**), no correlation reached statistical significance (*p* > 0.05). Repeating the analysis across all constructs including later described optoDRD1 variants (n = 15) correlation reached statistical significance for four parameters, three parameters showed a significant positive correlation with cAMP amplitude (receptor pLDDT, contact probability, and iPTM), while PAE had a negative correlation (*p* < 0.05; **Fig. S6b-c**). When the optoDRD1 constructs (c1–c5 and DRD1; n = 6) were analysed separately (**Fig. S8b-d**), the same parameters had significant correlations with higher rho values (ρ = 0.94, *p* = 0.005 for each of them). Qualitatively, the metrics separated the constructs into low-confidence models (JSR1-WT and JSR1-S199F), intermediate-confidence models (most chimeras) and the highest-confidence models (Gs-coupled receptors), although the wild-type Gs-coupled receptors also showed a variation of confidence metrics (**Table S3**). Given the small number of constructs, we regard these metrics as a filter for structural plausibility and not as quantitative predictors of efficacy.

JSR1-WT activation propagates through conserved Class A motifs and most chimeric receptors preserve key activation motifs including JSR1-DRD1 (**Fig. S7**). Notably, the non-functional JSR1-LSHR chimera lacks several of these conserved elements, highlighting their importance for the chimera function.

### Systematic optimization of the JSR1-S199F-DRD1 chimera

Since the JSR1-DRD1 chimera produced the strongest cAMP response, we adapted its design to the photoswitchable JSR1-S199F mutant. To optimise the construct and dissect the contribution of individual grafted regions, we systematically tested combinations of ICL2, ICL3, helix 8, and the CT from DRD1, generating five chimera variants (c1-5) each incorporating additional elements (**Fig. 3a**).

**Figure 3.**
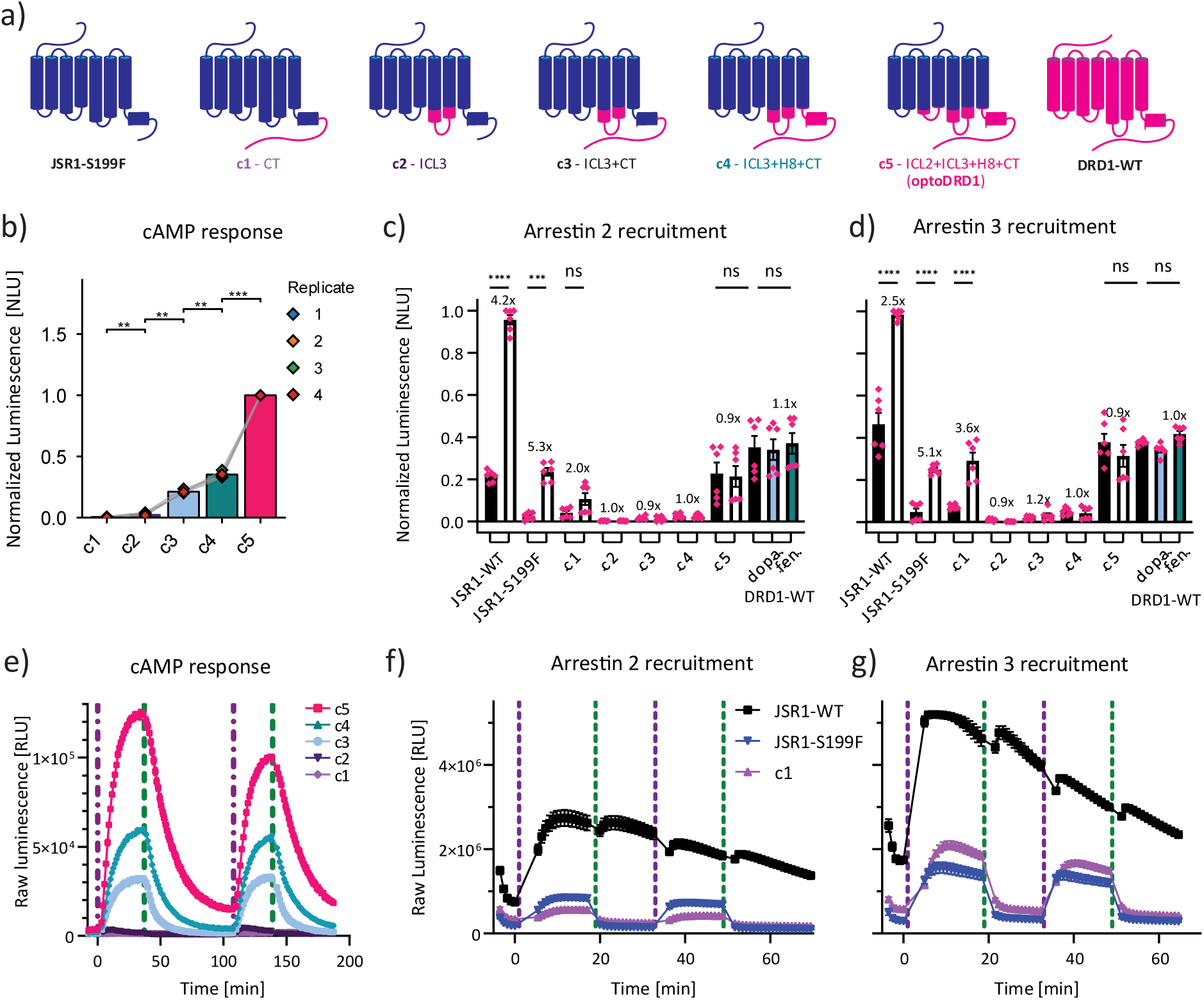
GloSensor cAMP responses and indirect arrestin recruitment to the cell membrane of different JSR1-S199F-DRD1 chimeras in HEK293T cells. **a)** Graphical depiction of different JSR1-S199F-DRD1 chimera designs (c1-c5; blue: JSR1, pink: DRD1). **b)** cAMP peak amplitudes of c1 to c5 normalised to the highest amplitude per biological replicate (n=4, colour-coded by replicate, bars show mean ± SEM. Pairwise comparisons were done by two-sided paired t-tests with Holm adjustment; brackets indicate significant pairs (** *p* < 0.01, *** *p* < 0.001). **c-d)** Indirect arrestin recruitment assay. Black columns indicate unstimulated conditions, and white columns were stimulated with 10^16^ photons cm^-2^, while DRD1 was stimulated with 10 µM dopamine (dopa.) and fenoldopam (fen.) as indicated, (\*\*\*\**p* < 0.0001, \*\*\**p* < 0.001, ns > 0.05). The numbers above the columns indicate the fold change compared to the unstimulated conditions. **e-g)** Time-resolved raw signalling data of different chimeras supplemented with 9-*cis*-retinal and illuminated with 10^16^ photons cm^-2^ at T = 0. The dotted lines indicate activation with violet light (385 nm) and deactivation with green light (525 nm). **e)** GloSensor cAMP assay of representative time course from replicate 3 in panel b, n = 2, mean ± SEM. Additional parameters including basal activity, fold change, 2^nd^ peak amplitude, raising and falling rates of all four replicates are displayed in Fig. S9. **f-g)** Indirect arrestin recruitment assay of representative dataset, n = 3, mean ± SEM. Only receptors with high stimulation-dependent arrestin recruitment are shown. JSR1-S199F was illuminated as indicated before and JSR1-WT was illuminated alternating with 525 nm and 595 nm. Time-resolved data of all individual chimeras are presented in Fig. S11.

Exchanging only the CT (c1) does not alter the signalling profile of JSR1-S199F, as violet light activation of this chimera leads to Gi-induced decrease of cAMP **(Fig. S8a)**. Exchanging only ICL3 (c2), increases cAMP response, in agreement with a previous report^35^. Combining of ICL3 and the C-terminus (c3) produces a 14-fold amplitude increase in Gs signalling compared to c2 (**Fig. 3b**). Adding ICL4 and helix 8 to c3 (generating c4) further enhances the Gs response. Including ICL2 in the final variant (c5) increases the absolute amplitude approximately two-fold relative to c4 (**Fig. 3b**). When comparing fold induction over baseline c3-c5 are not significantly different (∼30-fold increase to baseline; **Fig. S9c)**, because c5 also displays a correspondingly higher pre-stimulation baseline **(Fig. S9a, Table S5)**. ICL2 therefore raises the absolute output but not the dynamic range. Structural studies of DRD1 have shown that ICL2 establishes hydrophobic interactions with the Gαs protein^45^ and serves as binding site of positive allosteric modulators^46,47^. Our structural predictions support these experimental findings. Particularly, AF modelling of the c1-c5 variants with miniGs shows that c5 achieves the highest confidence scores at the receptor-G protein interface **(Fig. S8, Table S4)**, consistent with its highest cAMP amplitude.

To verify that the increased amplitude of cAMP signal of c5 stemmed from enhanced signalling capacity rather than elevated expression, we performed Western blot analysis. Expression levels did not significantly differ across variants. It is however worth to mention that we have not verified equal surface expression (**Fig. S10)**.

Beyond amplitude effects, we also examined how these structural elements influence cAMP signalling kinetics. This analysis revealed distinct patterns among the constructs. The ICL3 substitution alone (c2) decays after reaching its peak around 11 minutes regardless of green light deactivation **(Fig. S8a)**. A sustained cAMP activity is only achieved with a combined substitution of ICL3 and the C-terminus, highlighting the importance of these regions in maintaining sustained signalling kinetics. The activation rate constants are very similar among c3-5. However, the deactivation rates are slightly faster for c3 compared to c4 and c5 **(Fig. S9g-i, Table S5)**.

Since arrestins play a crucial role in GPCR desensitization, internalization and the initiation of arrestin-mediated signalling cascades, we assessed the recruitment of arrestin 2 and arrestin 3 to the cell membrane upon activation of JSR1-WT, JSR1-S199F, the JSR1-DRD1-chimera variants c1-c5 and DRD1. JSR1-WT indirectly recruits both arrestin 2 and arrestin 3 to the membrane upon light activation (**Fig. 3c-d**). Importantly, in the JSR1-S199F mutant this recruitment can be rapidly reversed by a deactivating light pulse, demonstrating that receptor deactivation is sufficient to disengage arrestin and potentially restore receptor responsiveness (**Fig. 3f-g**). Systematic replacement of JSR1 intracellular regions with their DRD1 counterparts revealed distinct contributions of each domain to indirect arrestin recruitment (**Fig. 3c-d**). Replacing the C-terminus alone (c1) reduces arrestin 2 recruitment while minimally affecting arrestin 3, suggesting isoform-selective interactions at the C-terminus. Replacing ICL3 (c2) almost completely abolished arrestin recruitment of both arrestin isoforms **(Fig. S11b**). This may indicate that ICL3 is either the dominant arrestin recruitment site of JSR1 or induces conformational changes, phosphorylation, or altered membrane abundance that could affect arrestin recruitment. Variants c3 and c4 show minimal light-induced arrestin recruitment, indicating that these substitutions alone are insufficient to confer DRD1-like arrestin engagement. In contrast, the addition of the ICL2 (c5) leads to a significant increase in basal arrestin recruitment (**Fig. 3c-d**), suggesting that ICL2 either provides an additional arrestin-binding interface or allosterically stabilises a receptor conformation with higher arrestin affinity. Notably, the arrestin recruitment profile of c5 most closely resembles that of DRD1, in line with our cAMP data. Both DRD1 and c5 display elevated basal arrestin association, and stimulation did not significantly increase recruitment above this baseline. This elevated basal recruitment likely reflects a high-affinity arrestin interaction in the absence of agonist, which may limit the dynamic range of light- or ligand-induced responses.

The chimera c5, which contains ICL2, ICL3, helix 8 and the C-terminus of DRD1, most closely resembles DRD1 in cAMP basal activity, cAMP signal amplitude **(Table S5)** and arrestin recruitment (**Fig. 3c-d**). This design is identical to the JSR1-DRD1 construct (**Fig. 2**), except that it is based on the photoswitchable JSR1-S199F mutant instead of the JSR1-WT. Because of the similarity to DRD1 and the high cAMP signal amplitude, we designated JSR1-S199F-DRD1 c5 as optoDRD1 and characterised it further. The chimeras c3 and c4 reach a lower cAMP amplitude, but they may be preferable for applications that require lower basal activity, faster shut-off or light-induced arrestin 3 recruitment.

Next, we tested which retinal isomer is required for a functional optoDRD1. 9-*cis*-retinal provided optimal starting conditions, producing an inactive receptor state that can be activated by violet light and deactivated with green light repeatedly. In contrast, 11-*cis*- or all-*trans*-retinal supplementation resulted in a partially or fully active state **(Fig. S12).** Nonetheless, following an initial green light deactivation pulse, the receptors reconstituted with these isomers display the same properties as the 9-*cis*-retinal reconstituted optoDRD1, and can be repeatedly switched on and off with violet and green light **(Fig. S12a)**. Time-course analysis of retinal supplementation (10 µM) revealed isomer-dependent gradually increasing cAMP levels over time, which is highest for all-*trans*-retinal but also present in 11-*cis*- and to a small extent in 9-*cis*-retinal **(Fig. S12b)**. Removing retinal from the assay buffer instead stabilizes the basal activity in dark conditions and leads to more complete deactivation (**Fig. 4f, S14a, c)**. We propose that, rather than binding 11-*cis*-retinal directly, optoDRD1 preferentially incorporates all-*trans*-retinal generated by spontaneous thermal isomerisation of 11-*cis*-retinal in aqueous solution at room temperature. This interpretation is supported by the observation that initial illumination at both 385 nm and 525 nm reduces cAMP in HEK cells supplemented with 11-*cis*- and all-*trans*-retinal, consistent with deactivation of a pre-existing active state bound to all-*trans*-retinal. In contrast, cells supplemented with 9-cis-retinal showed the expected activity increase upon 385 nm stimulation **(Fig. S12c-e)**. Molecular dynamics simulations by Costa et al. of JSR1-WT containing 9-*cis*-, 11-*cis*-, and all-*trans*-retinal revealed isomer-dependent differences in the long-range allosteric communication through helix 6 toward the intracellular G-protein-binding site. These findings indicate that the retinal-binding pocket is allosterically coupled to the G protein-binding region and may sense subtle differences between 9-*cis*- and 11-*cis*-retinal^48^. The apparent ability of optoDRD1 to incorporate all-*trans*-retinal may be advantageous for *in vivo* applications, as all-*trans*-retinal is present in many tissues, whereas the synthesis of 11-*cis*-retinal is restricted to the retinal pigment epithelium cells.

**Figure 4.**
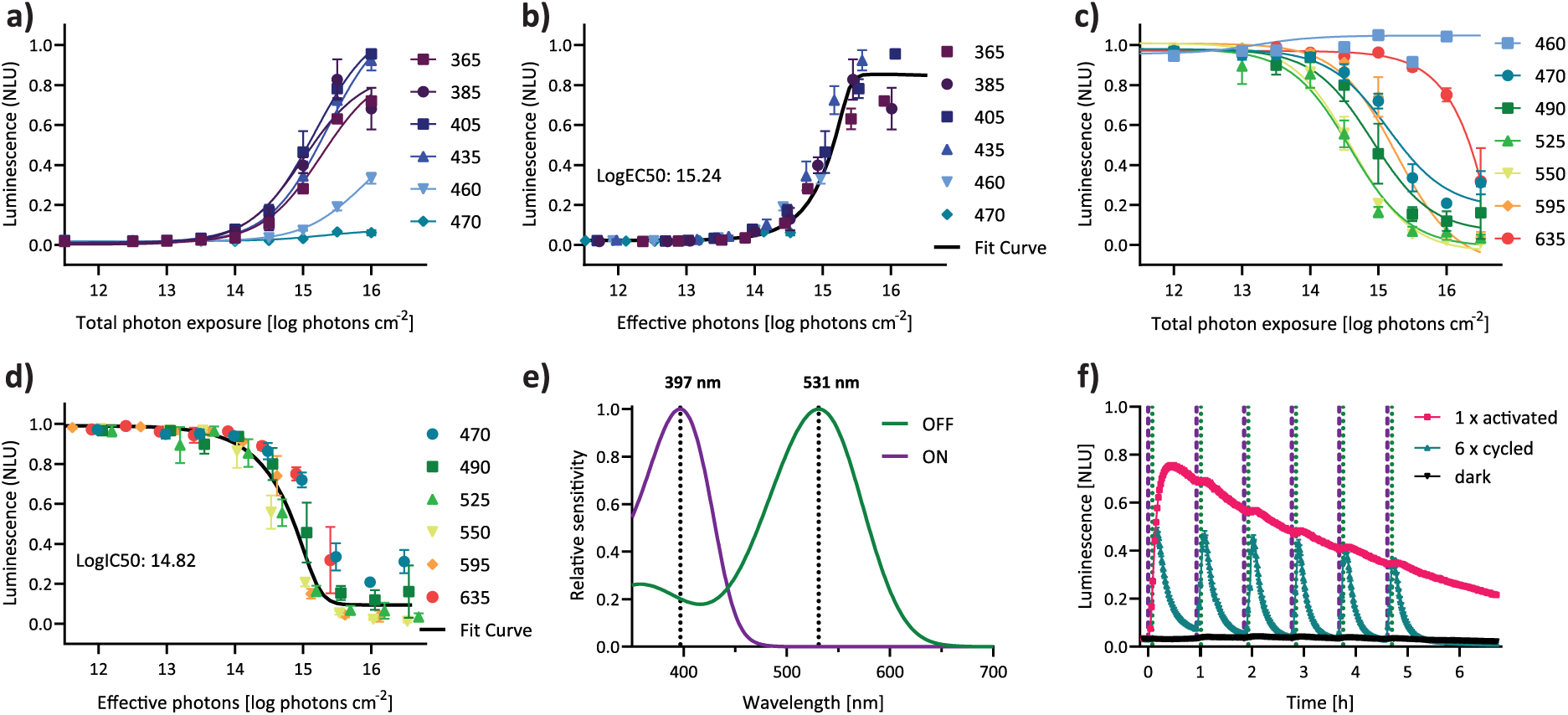
Light sensitivity of optoDRD1 measured by a spectral sensitivity assay in HEK293T cells. **a)** Dose-response curves for 1 s receptor illumination across intensities ranging from 10^11^ to 10^16.5^ photons cm^-2^ at the specified wavelengths for activation, n = 3 biological replicates. **(b)** Normalised dose response curves (from a) to effective photon flux, calculated based on the determined λ_max_ 397 nm. **c)** Dose-response curves for 1 s receptor illumination across intensities ranging from 10^11^ to 10^16.5^ photons cm^-2^ at the specified wavelengths for deactivation, n = 3 biological replicates. **(d)** Normalised dose response curves (from c) to effective photon flux, calculated based on the determined λ_max_ 531 nm. **e)** Govardovskii nomogram fit of signalling responses across six different wavelengths at ten different intensities each as shown in a-d, n = 3 biological replicates. The resulting λ_max_ values are 397 ± 1.08 nm for activation and 531 ± 1.25 nm for deactivation. **f)** Repeated stimulation of optoDRD1 in HEK293T cells with retinal-limited conditions (retinal was supplied only during overnight incubation, but not in the assay buffer). OptoDRD1 was activated with violet light and deactivated five minutes later with green light. This pattern was repeated six times. Values were normalised to the highest luminescence value of each biological replicate in the dataset, including both the JellyOp and optoDRD1 values of 4f and Fig. S14b-d.

Having established optoDRD1 as our lead construct, we next characterised its spectral sensitivity and signalling properties, comparing them to the native DRD1.

### Spectral sensitivity and bistable properties of optoDRD1

To establish the wavelengths and intensities required to control optoDRD1, we measured its spectral sensitivity using heterologous action spectra^49^ (**Fig. 4**). HEK293T cells expressing either JSR1-S199F with the Ca^2+^ indicator Aequorin, or optoDRD1 with the cAMP indicator GloSensor, were stimulated with varying light intensities at six wavelengths to generate irradiance response curves.

Ground-state activation of JSR1-S199F yielded a λ_max_ of 389 ± 0.54 nm (**Fig. S13**), consistent with the absorbance maximum of purified protein (λ_max_ = 385 nm), confirming that the spectral properties are preserved in the cellular context. OptoDRD1 showed a slightly red-shifted ground-state activation λ_max_ of 397 ± 1.08 nm (**Fig. 4a, b, e**), likely reflecting subtle changes in the retinal binding pocket environment introduced by the chimeric intracellular regions. The optoDRD1 active-state deactivation spectrum yielded a λ_max_ of 531 ± 1.25 nm (**Fig. 4c, d, e**). Notably, illumination at 635 nm could also deactivate optoDRD1, albeit requiring higher light intensities consistent with stimulation at the tail of the absorption spectrum. This extends the range of compatible light sources for experimental applications.

With approximately 130 nm separation between activation and deactivation peaks, both transitions reached saturation at approximately 10^16^ photons cm^-2^ s^-1^ (∼5 mW cm^-2^) for 1 s illumination pulses, demonstrating that optoDRD1 can be bidirectionally controlled at physiologically achievable light intensities with minimal spectral crosstalk between the two control wavelengths.

To demonstrate the advantages of using a bistable opsin as a scaffold for the optoGPCRs we measured cAMP signalling without added retinal in the assay buffer and repeatedly switched the receptor on and off six times over a 6 h period. Activating the receptor once maintained elevated cAMP levels over the full 6 h, while repeated 5-minute activation periods (activating the receptor and deactivating it 5 minutes later) produced reproducible signal spikes with an amplitude of ∼60% of the amplitude achieved with sustained activation, with a modest decline to ∼50% by the sixth cycle (corresponding to a decrease to ∼73% of the first-cycle spike amplitude), demonstrating that optoDRD1 supports both sustained and repeatable, on-demand activation over multiple light cycles (**Fig. 4f**).

### Temporal signalling dynamics and G protein selectivity of optoDRD1

To determine whether light-activated optoDRD1 reproduces the signalling profile of DRD1, we compared the temporal dynamics of cAMP responses and G protein selectivity profiles between the two receptors. We first examined the temporal dynamics of cAMP responses, comparing optoDRD1 to DRD1 activated with the DRD1-specific agonist fenoldopam and deactivated with the antagonist SKF83566. Both receptors showed comparable peak amplitudes and sustained signalling over hours, with apparent cAMP-response rate constants that were not significantly different between optoDRD1 and DRD1 (Fig. 5a-b, Table S5; *p* > 0.57). The cAMP signal integrates several steps: receptor activation, Gs coupling, adenylyl cyclase activity, cAMP degradation by phosphodiesterases (PDEs) and the response time of the biosensor. The apparent kinetics therefore reflect this integrated signal rather than the intrinsic kinetics of receptor activation, which occur on much faster (sub-second to second) timescales^50^.

**Figure 5.**
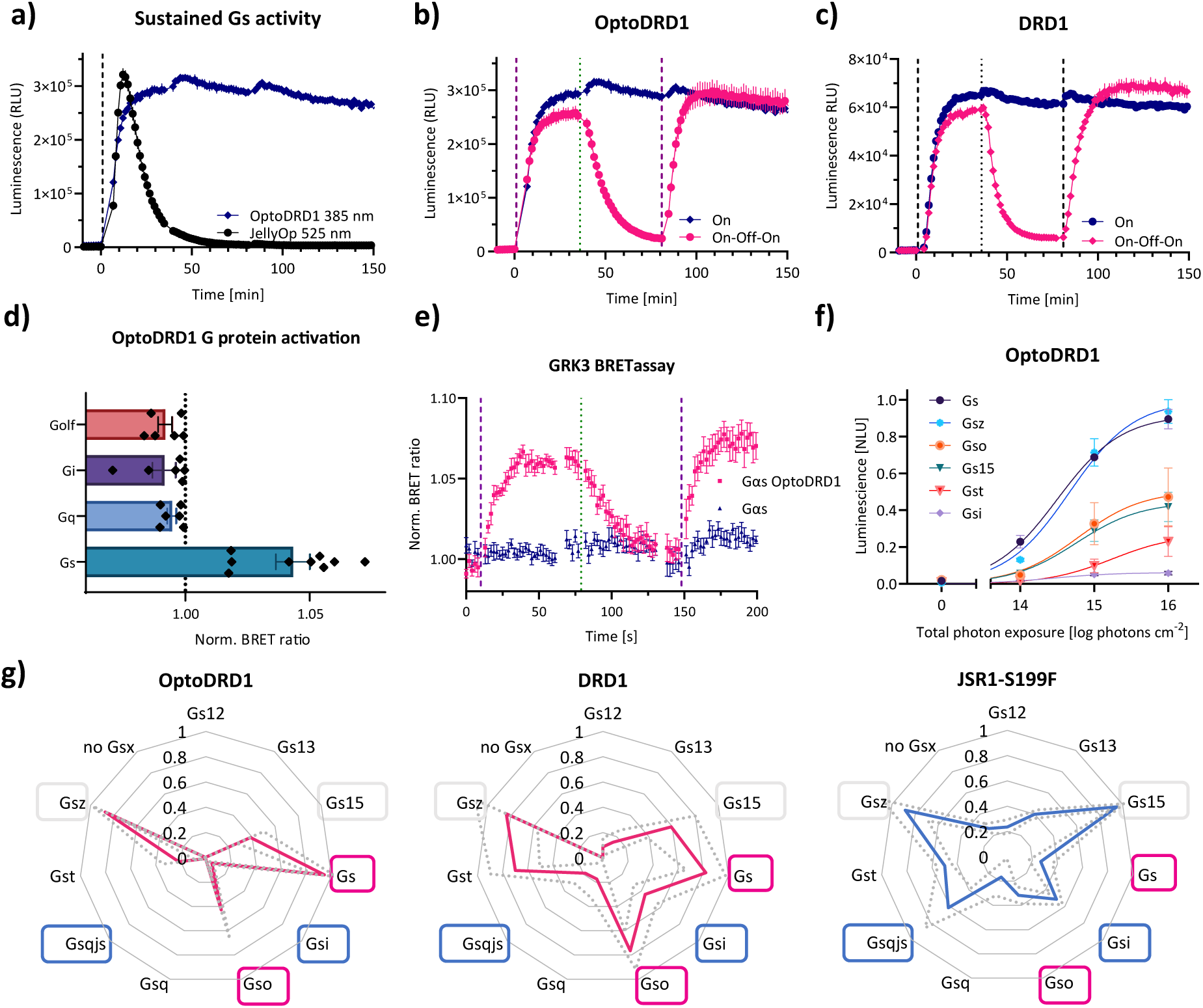
G protein selectivity and kinetics of optoDRD1. Comparison of optoDRD1, DRD1, and JellyOp signalling profiles with the GloSensor cAMP assay (a-c) in HEK293T cells. **a)** Response curves of optoDRD1 and JellyOp reconstituted with 9-*cis*-retinal stimulated once at the dashed line for 1 s (10^16^ photons cm^-2^). **b)** Response curves of optoDRD1: the dark blue curve (‘On’) shows single activation with 385 nm. The pink curve (‘On-Off-On’) corresponds to sequential stimulation with 385 nm, 525 nm and 385 nm light (all at 10^16^ photons cm^-2^). **c)** Response curves of DRD1: the dark blue curve (‘On’) shows the effect of 10 nM fenoldopam (agonist). The pink curve (‘On-Off-On’) shows the effect of additional treatment (T = 36 min) with 100 nM SKF83566 (antagonist), followed by addition of 1 µM fenoldopam (T = 81 min). For **a-c**, traces represent mean ± SEM of n = 3 from a representative dataset measured under matching experimental conditions. Quantitative comparison of n = 3 biological replicates is in Table S5. **d)** OptoDRD1 G protein selectivity from BRET G protein activation assay in HEK293T cells, displaying mean ± SEM, n = 6-9. **e)** OptoDRD1 in a BRET G protein activation assay in HEK293T cells, showing Gs activity after illumination for 1 s with 385 nm (10^16^ photons cm^-2^), 525 nm (10^16^ photons cm^-2^), and 385 nm (10^16^ photons cm^-2^). The data represent mean ± SEM, n = 6. **f)** Dose response curve of optoDRD1 of the Gsx assay with HEK293S ΔGs cells depicting G proteins that had a response of at least 0.1 of the normalised maximal response. **g)** Summary of Gsx assay in HEK293S ΔGs KO cells for the indicated receptors at 10^16^ photons cm^-2^ or 10 nM fenoldopam for DRD1. Response curves were normalised by subtracting the unstimulated condition, subsequently the average of five values after stimulation reached maximum was calculated. These values for each receptor were then normalised from 0 to 1. Data represent mean ± SEM (grey dotted line) n = 3.

For comparison, we tested JellyOp, a commonly used Gs coupled opsin^22^. When optoDRD1 and JellyOp were compared under matched conditions (same plate and cell batch, 2 µM 9-*cis*-retinal, identical DNA amounts, same promoter), both reached a similar peak amplitude (Fig. 5a). However, we did not independently verify equal surface expression of optoDRD1 and JellyOp, so differences in absolute amplitude should be interpreted with caution. The two receptors differed instead in the persistence of the signal and in repeated switching (Fig. 5a, S14). JellyOp produced a robust but transient response, rising and then spontaneously decaying with a peak at ∼ 11 minutes without reaching a sustained plateau consistent with previous reports that JellyOp does not require a deactivating light stimulus (**Fig. 5a**). This contrasts with the light-controlled bistability of optoDRD1, which remains in a stable active state until deliberately deactivated by green light. The extracted rate constants for conditions where receptor was only activated once without any deactivation indicate a much faster deactivation rate for JellyOp (0.09 min⁻¹) than for DRD1 (0.001 min⁻¹) and optoDRD1 (0.003 min⁻¹), which both decay very slowly. In the 6h measurement optoDRD1 has a half-life (t_1/2_) of about 5h (Fig. 4f). The spontaneous decay of JellyOp (0.09 min⁻¹) is in the same order of magnitude to the light-induced deactivation of optoDRD1 (0.05 min⁻¹; **Fig. 5c, Table S5**). Because both receptors share the same reporter system, this suggests comparable cAMP clearance kinetics once deactivation begins, whether spontaneous (JellyOp) or light-triggered (optoDRD1). Together, these results indicate that the chimeric design of optoDRD1 enables sustained, light-controlled activation similar to DRD1 cAMP responses, when continuously exposed to the agonist. JellyOp may be the preferred choice for applications requiring short pulses of activation without requiring control over the length of the signalling active state.

We next assessed the G protein selectivity profile of optoDRD1 using a BRET-based G protein activation assay with native human G proteins. This assay revealed activity exclusively for Gs, with no detectable coupling to Gq or Gi (Fig. 5d,e), consistent with the absence of Gq activation in the orthogonal Aequorin assay (Fig. S15). These results demonstrate that optoDRD1 selectively activates native Gs, recapitulating the primary coupling preference of DRD1^31^. To complement this analysis, we additionally employed the Gsx assay^51^ in HEK293S ΔGs cells lacking endogenous Gαs protein^52^. The Gsx assay probes G-protein selectivity via chimeric Gα proteins consisting of a Gs core, which directs signalling through cAMP increases, with variable C-terminal sequences for each of the Gα subtypes. This provides a proxy of G-protein selectivity, as the majority of the GPRC-Gα binding interface is determined via the Gα C-terminal tail. In this assay, both DRD1 (**Fig. 5g, middle)** and OptoDRD1 (**Fig. 5g, left)** showed similar Gsx preferences with the highest activity for Gs, followed by Gsz, and weaker activity for Gso, Gst and Gs15. In contrast, the control construct JSR1-S199F showed weak activation across Gsx chimeras, with the relatively highest activity for Gsi and Gsqjs (jumping spider Gq C-terminus; Fig. 5g, right), neither of which was activated by optoDRD1. One important caveat with interpretation of this dataset is that in at least 50% of published DRD1-Gαs structures DRD1 also makes contact with the Gαs core (Table S6). GPCR-Gsx selectivity mediated primarily by the C-terminal tail may therefore not provide a true assessment of G protein selectivity. Together, the BRET data show that optoDRD1 selectively activates native Gs (Fig. 5d-e), and the Gsx assay shows that its C-terminal preference resembles that of DRD1.

### OptoDRD1 in different cell types

To evaluate the broader applicability of optoDRD1 beyond HEK293 cells, we tested its function in pancreatic β-cells and a neuronal cell line, two contexts in which Gs-mediated cAMP signalling plays well-established physiological roles. We first examined whether optoDRD1 could modulate insulin secretion in EndoC-βH5® human β-cells, a cell line closely resembling native human pancreatic β-cells in physiology and function^53^. Gs-coupled receptors including GLP-1R and GIPR are known to potentiate glucose-stimulated insulin secretion (GSIS) through cAMP-dependent mechanisms^26^ (**Fig. 6a**). Light stimulation of optoDRD1 reconstituted with 9-*cis*-retinal increased insulin release 1.5-fold compared to unstimulated controls (**Fig. 6b**). Insulin release was elevated to a similar extent in cells reconstituted with all-*trans*-retinal in the absence of light stimulation, consistent with the agonist activity of all-*trans*-retinal observed in HEK293T cells **(Fig. S12)**. Since the experimental conditions did not allow to extract released insulin from the same well after activation and after deactivation, we started from an active receptor by supplementing all-trans-retinal and to observe the effect of deactivation with green light. Green light applied immediately prior to glucose stimulation did not reduce insulin secretion in all-*trans*-retinal-reconstituted cells relative to the non-deactivated condition (**Fig. 6b**). This may reflect the slow decay of cAMP compared to the rapid first-phase insulin response, which peaks within the first five minutes of glucose exposure^54^, such that cAMP levels remained elevated despite receptor deactivation.

**Figure 6.**
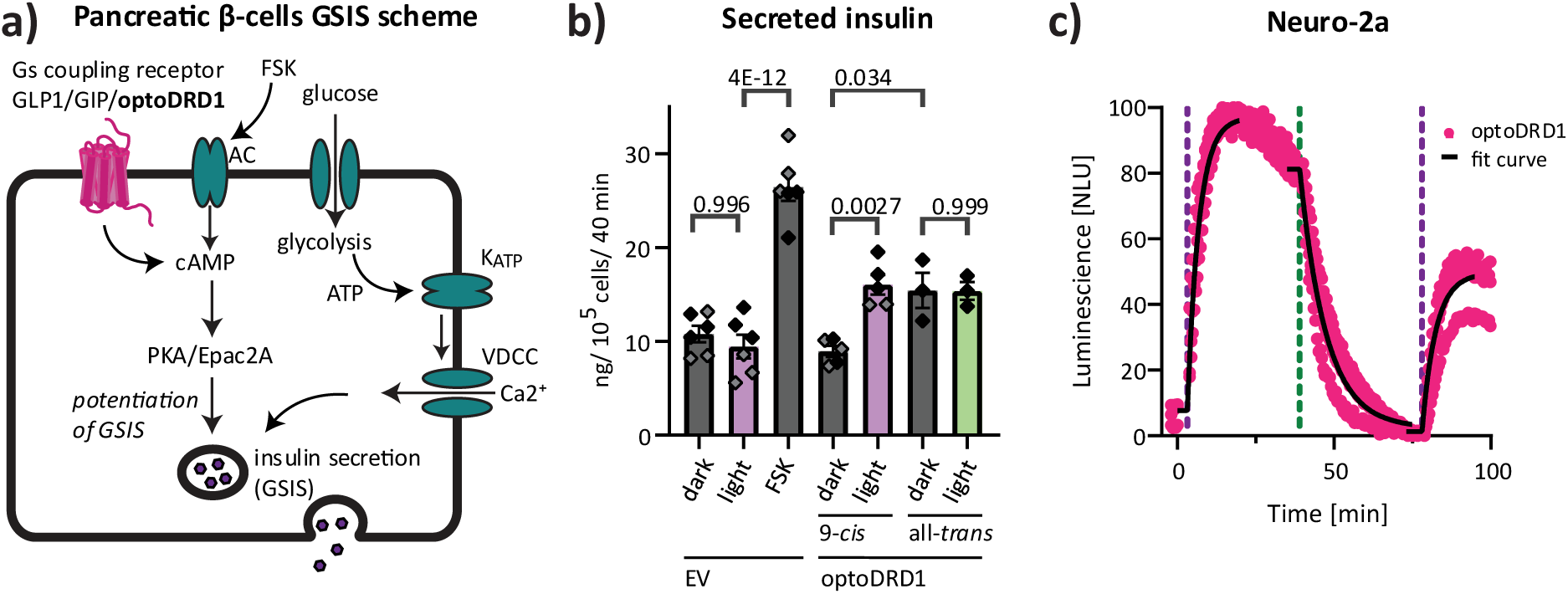
Activity of optoDRD1 in other cell types. **a)** Schematic showing the mechanism of glucose-dependent insulin release in pancreatic β-cells, which is amplified via Gs signalling (adenylyl cyclase (AC), protein kinase A (PKA), K_ATP_ channel (K_ATP_), voltage-dependent calcium channel (VDCC)). **b)** OptoDRD1 activation enhances insulin release from human pancreatic β-cells (EndoC-βH5®) upon glucose stimulation. All conditions were stimulated with 11 mM glucose, with forskolin (FSK) as positive control and light (violet or green) as indicated by the column colour (empty vector (EV); the numbers indicate the adjusted *p*-values). Statistical significance was determined by one-way ANOVA followed by Tukey’s HSD test for multiple comparisons. Diamonds represent individual measurements; bars show the mean ± SEM, n = 3-6 per group (replicates of week 1, black; week 2, grey). **c)** In Neuro-2a cells, optoDRD1-mediated cAMP responses to light activation and deactivation were monitored using the GloSensor cAMP assay. Cells were incubated with 2 µM 9-*cis*-retinal and illuminated at T = 2 min and T = 79 min with a 385 nm light pulse and at T = 40 min with a 525 nm light pulse of 1 s (10^16^ photons cm^-2^). Data represent mean ± SEM of n = 3. Black lines represent the fitted curve.

We next examined whether optoDRD1 functions comparably to HEK293T cells in a neuronal context by measuring cAMP levels in Neuro-2a cells. OptoDRD1 reversibly activated cAMP signalling upon stimulation with violet light and deactivation with green light similar to HEK293T cells (**Fig. 6c, Table S5)**. However, the effect of 11-*cis*-retinal on baseline receptor activity differed between cell types. In HEK293T cells, 11-*cis*-retinal activated the receptor in the dark, necessitating green-light deactivation before a violet photoactivation response could be detected. In Neuro-2a cells, 11-*cis*-retinal produced only partial receptor activation, leading to further activation with 385 nm illumination and deactivation below initial cAMP levels with 525 nm illumination **(Fig. S12f-h)**. This difference does not impair the utility of optoDRD1 for light-controlled cAMP modulation in this cell type, but its mechanistic basis warrants further investigation and may reflect cell-type-specific differences in retinal metabolism.

## Discussion

In this study, we have developed optoDRD1, a bistable optoGPCR that recreates the signalling profile of DRD1 and enables bidirectional control of Gs signalling with light. Despite the major advantage of bistable photoswitchable opsins being light-controlled deactivation, most engineered optoGPCRs currently use monostable opsins that photobleach after activation^34^. We therefore selected the bistable and photoswitchable JSR1-S199F point mutant as opsin scaffold for chimera generation^14^. JSR1 has available structures of both the ground and active states, making it an ideal opsin for rational engineering^12,19^. Our optoDRD1 design provides several advantages over existing monostable opsin-based Gs-signalling optoGPCRs. Specifically, (i) it does not photobleach, (ii) it enables reversible control, and (iii) it can form functional photopigment using all-*trans*-retinal, which is widely available throughout the body, while 11-*cis*-retinal has a limited tissue distribution and is mainly present in the retina^55,56^.

OptoDRD1 activates with violet light (λ_max_ = 397 nm) and deactivates with green light (λ_max_ = 531 nm) at high sensitivity (EC50 ≈ 10^15^ photons cm^-2^ s^-1^, ∼0.5 mW cm^-2^), approximately 100-fold more sensitive than widely used microbial opsins such as ChR2^57,58^, a sensitivity range compatible with established *in vivo* optogenetic paradigms. The shorter wavelengths required for optoDRD1 activation penetrate tissue less effectively than red light and may carry greater phototoxicity risks, representing a practical limitation for deep-tissue applications. Recent advances in two-photon activation have demonstrated that deep tissue stimulation of UV-sensitive opsins is feasible^6,59^. OptoDRD1 may therefore be suitable for two-photon activation, offering deeper tissue penetration and single-cell spatial resolution within living organisms^60^.

To date, most *in vivo* applications of bistable photoswitchable optoGPCRs have used Gi- or Gq-coupled opsins^6–8,15^. JellyOp remains the most widely used Gs-coupled optoGPCR and has been applied to photostimulate Gs signalling in the heart^61^. It can be activated repeatedly and rapidly with light, and it deactivates spontaneously with its own intrinsic kinetics, with the cAMP signal starting to decay within 15 minutes in cellular assays^22,62^ (Fig. 5a, S14). This transient response can be advantageous when brief, Gs signalling is desired. However, sustained modulation of downstream pathways requires repeated light stimulation. On the other side, shutting JellyOp down by UV light before its spontaneous decay leads to desensitisation that prevents re-activation for several minutes^22^. OptoDRD1 offers a complementary kinetic profile. It maintains Gs activity for several hours, closely matching the cAMP kinetic profile of ligand-activated DRD1 (Fig. 5, Table S5). The cAMP kinetics reported here reflect the integrated receptor, G protein, adenylyl cyclase, PDE and reporter system and are therefore apparent cAMP-response kinetics rather than intrinsic receptor kinetics. Moreover, optoDRD1 can be activated and deactivated with light at defined time points, which enables temporal dissection of receptor-mediated Gs signalling. Under the same experimental conditions, both JellyOp and optoDRD1 reached a similar peak amplitude (Fig. 5a, Table S5), whereas the two differed in signal persistence, fold-change and repeated responses under retinal-limited conditions (Fig. S14). Receptor surface expression could not be reliably quantified between optoDRD1 and JellyOp. Therefore, we focused on response persistence, light-dependent termination and repeated switching, because these measures are less sensitive to expression differences than absolute amplitude.

The prolonged activity of optoDRD1 is consistent with the absence of arrestin recruitment to DRD1 and optoDRD1 upon stimulation. This suggests that arrestin-mediated desensitisation contributes minimally to signal termination under our experimental conditions. The elevated basal arrestin recruitment present in both, DRD1 and optoDRD1 may reduce the detectable dynamic range of agonist-induced recruitment. Discrepancies with previous reports, which showed arrestin recruitment to DRD1 following agonist stimulation, may therefore reflect differences in experimental conditions and/or receptor expression levels^63,64^. Since our arrestin readout is an indirect assay, we cannot exclude low level or transient recruitment, and the absence of a detectable signal should not be interpreted as an absence of arrestin engagement.

To demonstrate the proof-of-principle of optoDRD1, we examined its ability to modulate endogenous Gs-mediated processes. We demonstrate that activated optoDRD1 augments glucose-stimulated insulin secretion in human primary-like pancreatic β cells (EndoC-βH5®). This is possible both through optoDRD1 reconstituted with 9-*cis*-retinal and activated with violet-light, as well as optoDRD1 directly supplemented with all-*trans*-retinal (**Fig. 6**). Although violet light activation immediately prior to glucose stimulation was sufficient to increase insulin release, deactivation directly before did not significantly reduce insulin release, likely reflecting the relatively slow decay of cAMP compared to the rapid first-phase insulin response, which peaks within the first five minutes of glucose exposure^54^. These results confirm that all-trans-retinal supplementation is sufficient to activate optoDRD1. This is beneficial for tissues lacking 11-cis-retinal, but it comes at a cost: all-trans-retinal reconstitution produces a receptor population with basal Gs activity before any light stimulation (Fig. S12). This activity can be reduced by a brief green-light pre-illumination that returns the receptor to its inactive ground state, or by reconstituting with 9-cis-retinal. Prolonged high Gs activity might induce stress responses in some cells^65,66^, other cells might be able to adapt or even benefit from it^67,68^. In EndoC-βH5 cells, overnight incubation with all-*trans*-retinal increased insulin release to a similar extent as light-activated 9-*cis*-retinal-reconstituted optoDRD1 (Fig. 6b), indicating that these cells remained responsive after prolonged exposure to all-*trans*-retinal. However, we did not measure intracellular cAMP or receptor surface levels in these cells, and we cannot exclude adaptation, desensitisation or altered expression. Two points need to be tested for each new cell type or tissue. First, it must be determined whether endogenous retinal is sufficient to reconstitute optoDRD1 or whether it must be supplemented. Second, it must be determined whether optoDRD1 raises Gs activity in the dark and whether cells tolerate this. If basal activity is a problem, it can be reduced by using 9-cis-retinal or green-light illumination. However, optoDRD1 may still be unsuitable for applications that require a low dark-state signal.

Beyond its utility as an engineering scaffold, JSR1-S199F has signalling properties of independent interest. JSR1 has primarily been characterised as a Gq-coupled receptor, but we show that it also signals via Gi in human cells and recruits arrestin 2 and arrestin 3 efficiently, in a light-dependent and reversible manner. Analysis of the chimeras showed that ICL3 of JSR1 is required for arrestin 3 recruitment, whereas replacing the C-terminus had only a minimal effect. This could mean that ICL3, rather than the C-terminus, is the primary arrestin-interacting site in JSR1, or that the DRD1 C-terminus can compensate for the JSR1 C-terminus because it serves a similar function. Within the flexible ICL3, a sequence motif (SxxSxED) containing two serines, a glutamate and an aspartate may contribute to the high arrestin recruitment, since arrestin-binding interfaces are typically enriched in negative charges, such as those provided by phosphoserines. However, replacing ICL3 could also alter receptor conformation, phosphorylation or membrane abundance, any of which could affect arrestin recruitment independently of this motif. Arrestin recruitment to JSR1-S199F is rapidly reversed by green light (**Fig. 3f-g**). As arrestins are involved not only in receptor desensitisation and internalisation but also in initiating signalling cascades, it will be important to determine whether light-induced arrestin recruitment triggers arrestin-dependent effects such as ERK1/2 activation. The bistability of JSR1-S199F is particularly advantageous in this context, because recruitment can be both induced and terminated by light, without the need of using antagonists that may not reach an internalised receptor. Together with its Gq and Gi coupling, JSR1-S199F may therefore serve as a promising scaffold for optogenetic dissection of arrestin-mediated signalling with high spatiotemporal control.

The altered signalling profile of JSR1-S199F, which shows a markedly enhanced Gq response and a modestly increased Gi response relative to JSR1-WT (**Fig. 1a**), may reflect changes in the conformational ensemble generated by the retinal-binding pocket. Ser199 forms part of the hydrogen-bond network linking the protonated retinal Schiff base to the E194 counterion, and molecular dynamics simulations indicate that replacing this residue with phenylalanine perturbs this network and the associated interactions involving Tyr126 and Tyr293^69,70^. Recent simulations further show that retinal configuration influences communication between this region, Trp290, TM6 and the intracellular transducer-binding surface^48,71^. S199F may therefore shift the relative populations of receptor conformations available for G-protein and arrestin engagement, favouring states that couple more efficiently to Gq, rather than simply changing overall receptor activity. Because the GRK3-BRET assay reports Gβγ release, the minor Gq response of JSR1-WT could reflect either weaker Gq coupling or reduced G protein turnover (for example, stable, slowly dissociating complexes); our data cannot distinguish between these possibilities. As the simulations did not include G proteins or arrestins, this interpretation remains a testable hypothesis rather than direct evidence of transducer-specific selectivity. Testing it would require simulations of receptor-transducer complexes.

One of the main challenges in optoGPCR engineering is reduced signalling efficiency compared to native receptors^34^, potentially reflecting both lower expression levels and less efficient G protein activation. Here, cAMP signalling amplitude was highly dependent on the parent Gs-coupling receptor. Of seven engineered chimeras based on Class A (rhodopsin-like) GPCRs, only the JSR1-DRD1 chimera displayed high signalling efficiency (**Fig. 2**). Consistent with this, AF analysis predicted that optoDRD1 has higher confidence scores at the receptor-G protein interface than the other chimeras. These scores correlated significantly with experimental signalling efficiency when all chimeras described in the manuscript were included **(Fig. S6)**, but not for the initial screening chimeras alone. Given the small number of constructs and the dominance of a few high-signalling complexes, we regard these metrics as a useful tool for guiding GPCR chimera design, checking structural plausibility and prioritizing candidates for experimental validation and not as quantitative predictors of efficacy.

In agreement with previous GPCR engineering studies^16,42,72^, our JSR1-S199F-DRD1 variants show that inclusion of ICL2, ICL3, helix 8 and the C-terminus is sufficient to reconstitute native-like cAMP signalling and DRD1-like arrestin recruitment. Each element contributes distinctly: ICL3 initiates Gs coupling and is associated with minimal assay-detectable arrestin recruitment, the C-terminus sustains signalling duration, helix 8 amplifies the cAMP response amplitude, and ICL2 provides further enhancements to both Gs coupling and basal arrestin engagement. Earlier chimeras carrying the JellyOp ICL3 established Gs coupling in bistable opsins and were mainly used to study opsin light sensitivities via a cAMP assay, but they gave modest responses and did not combine sustained Gs output with repeatable switching^35^. The advance of optoDRD1 is therefore not the redirection to Gs by an intracellular loop swap alone, but the combination of the JSR1-S199F spectral switch, systematic multi-domain optimisation, robust and prolonged Gs output, and repeatable bidirectional light control (**Fig. 5, S14)**.

DRD1 has emerged as a particularly suitable receptor for opsin chimeras. Previously, Morri et al. included DRD1 in their large screen of bovine rhodopsin optoGPCRs, simultaneously exchanging ICL1, ICL2, ICL3, and the C-terminus. This resulted in one of the largest cAMP-dependent signals among their constructs^72^. However, this optoDRD1 showed high basal activity, suggesting aberrant constitutive signalling^72^. A separate bovine rhodopsin-DRD1 chimera was also successfully employed in the mouse nucleus accumbens, mediating behavioural effects on social interactions^73^. More recently, Zhou et al. created an optoDRD1 using bovine rhodopsin and the DRD1 of *Drosophila melanogaster*^42^. Their optimised design retained the ICL1 of bovine rhodopsin and introduced additional DRD1-derived residues in helix 8 (7.56-8.49). This optoDRD1 matched the signalling profile of native *Drosophila* DRD1 and enabled optical control of locomotion and learning behaviour *in vivo*^42^. However, all these previous designs used monostable opsins as scaffolds. Our optoDRD1 advances on these designs by enabling bidirectional light-triggered control–both activation and deactivation–addressing the high basal activity reported in prior designs while maintaining robust signalling in both HEK293 and the neuroblastoma Neuro-2a cell line. The ability to deactivate signalling with green light at any time point provides an antagonist-like modality, enabling precise temporal manipulation of Gs pathways not previously achievable with existing opsins.

OptoGPCRs control cellular systems through GPCR signalling pathways rather than the ion channel conductance mediated by type I opsins, enabling spatiotemporally precise analysis of specific GPCR signalling pathways. While photoswitchable ligands, such as the reversibly membrane-anchored MP-D1_ago_ or photoswitchable serotonins^74,75^, in principle can achieve similar precision without the need to express a recombinant receptor, optoGPCRs offer distinct advantages: their expression can be targeted to specific cell types and tissues, they are unaffected by endogenous ligands or non-specific binding of photoswitchable ligands to off-target receptors, and light activation eliminates the need to administer drugs, reducing systemic off-target effects. Unlike Designer Receptors Exclusively Activated by Designer Drugs (DREADDs), which are commonly used to study GPCR activation in neuronal systems^76^, optoGPCRs provide more precise temporal control of activation and deactivation without requiring ligand administration, making them valuable tools for dissecting specific GPCR pathways^34^. The choice between ligand-based, photopharmacological, or optoGPCR approaches should therefore be guided by the specific experimental requirements of the study.

Together, these results establish optoDRD1 as a bistable, Gs-specific optoGPCR that combines spectral reversibility, high light sensitivity, and native-like cAMP signalling kinetics. Additionally, we show that JSR1-S199F strongly recruits arrestin 2 and 3 in a light-sensitive manner, and this recruitment can be rapidly reversed by deactivating the receptor with green light. The light-mediated control of Gs signalling provided by optoDRD1 may be particularly valuable for investigating dopaminergic neuronal pathways implicated in learning and memory, Parkinson’s disease models, and Gs-mediated peripheral organ functions such as pancreatic β cell insulin secretion. OptoDRD1 thus represents a valuable addition to the optoGPCR toolbox for dissecting Gs-mediated signalling with spatiotemporal precision.

## Materials and Methods

### cDNA constructs

The generated protein sequences used in this study are listed in supplementary data. The GPCR open reading frames were synthetised by TwistBioscience and cloned into the mammalian expression vector pcDNA3.1+ with KpnI and XbaI cleavage sites (Amino acid sequences in SI). Construct contains the open reading frames of the GPCRs and a C-terminal 1D4 epitope tag (TESTQVAPA) which is connected to the GPCR with a GS linker (BamHI cleavage site). The coding sequence for DRD1 is based on NM_000794.5. The JSR1 gene was expressed with an N-terminal SNAP-tag (SI) and codon optimised for homo sapiens. Site-directed mutagenesis of JSR1 genes and chimeras was carried out using Phusion® High-Fidelity PCR Master Mix with GC Buffer (NEB) to introduce S199F according to the manufacturer’s instructions. JellyOp construct (Addgene plasmid # 41432) was previously described^62^. For reporting cAMP levels with bioluminescence, the pGloSensor-22F cAMP plasmid (Promega, E2301) was used. The Gsx chimeras were used as previously described^51^. For measuring indirect arrestin recruitment the plasmids (AG10-11s-CAAX, AK1-114-bArrestin1, AK1-114-bArrestin2) were used as described previously^77^. For the BRET G protein activation assay, GRK3-nLuc, Gα subunits, Gβ-split Venus and Gγ-split Venus subunits were used as described previously^78^. To capture intracellular Ca^2+^ levels, the pcDNA5/FRT/TO mtAeq plasmid^79^ was used, which expresses the Ca^2+^ sensitive luminescent aequorin protein. All plasmids were verified by sequencing prior to use.

### Cell culture

HEK293T cells, HEK293S ΔGs cells^52^ or Neuro-2a cells, were cultured at 37°C in culture medium containing: Dulbecco’s modified Eagle’s medium (DMEM) 4.5 g L–1 D-glucose, sodium pyruvate and L-glutamine (BioConcept) with 10% fetal bovine serum (FBS; Sigma-Aldrich) and penicillin/streptomycin (PANBiotech, 100 U/mL and 0.01 mg/mL respectively) in a 5% CO_2_ atmosphere.

### Transfections

Cells were transiently transfected with Lipofectamine 2000 (Invitrogen), according to the manufacturer’s protocol. After 4-6 h the medium was replaced with fresh medium, supplemented with retinal if required. Retinal was added in dim red-light conditions (680 nm long-pass filter) and cells were strictly handled under these conditions thereafter. For transfections performed in 12- or 24-well format, cells were detached with Versene (biowest) or trypsin and redistributed into 96-well-plates at 100 µL per well.

### Immunocytostaining

Cells were seeded 6 h after transfection onto a poly-D-lysine (Thermo Fisher Scientific) coated glass cover slip at density of 1.25 x 10^5^ cells per well in a 24-well plate. 24 h after transfection cells were fixed with 4% paraformaldehyde (Sigma-Aldrich) in PBS and stained with 0.1 µg/mL DAPI staining for 10 minutes at room temperature. Cells were permeabilised for 5 minutes with 1% NP40 in PBS and then blocked with 1% BSA in TBST for 30 minutes. Cells were incubated for 1 h with 1:1000 diluted 1D4-Rho mouse antibody 3.5 mg/mL (Cell Essentials, Inc) in blocking buffer. After washing with PBS cells were incubated with 1:1000 Alexa Fluor 594 conjugated anti-mouse secondary antibody (MedChemExpress). Cells were mounted with Gelvatol on glass slides and stored in the dark. The sample was examined on a Leica Stellaris microscope. Images were taken under specific band pass filter sets and colour-combined images were used.

### Western Blot

HEK293T cells were seeded at density of 2.5 x10^5^ cells per well in 24-well plate 24 h before transfection. Cells were transfected with expression vectors of the relevant opsin 250 ng of total DNA per well. After 24 h, cells were detached with versene and pelleted with ice-cold PBS. Cells were incubated in PBS with Benzonase for 15 minutes and loaded with loading buffer onto a bio-rad mini-PROTEAN TGX gel, electrophoresed, and turbo plotted onto a membrane and blocked with 3% BSA in TBST for 1 h. Membranes were incubated for 1 h with 1:1000 diluted 1D4-Rho mouse antibody 3.5 mg/mL (Cell Essentials, Inc) in blocking buffer. After washing with TBST membranes were incubated with 1:1000 HRP-conjugated goat anti-mouse secondary antibody (Invitrogen).

### Purification of JSR1-S199F and measurements of UV-vis spectra

JSR1-S199F protein was expressed in Hi5 insect cells with Baculovirus for 48 h. The cell membranes were washed by ultracentrifugation and reconstituted with 9-*cis*-retinal overnight. Every step after retinal addition was carried out in dim red-light conditions. Membranes were solubilised in 1% LMNG, affinity purified with Strep-Tactin XT resin and eluted with 0.05 mg/mL homemade 3C HRV protease. The UV-vis spectra were recorded on an HPLC Infinity II Diode Array Detector WR (Cat. Number G7115A) (Agilent, California, USA) during an analytical Size Exclusion Chromatography (SEC) run. The purified receptors (10 µl) were injected at 20°C onto a Zenix-C SEC300 4.6×150 mm column (SEPAX Technologies, Delaware, USA) pre-equilibrated in SEC buffer (20 mM HEPES pH 6.5, 300 mM NaCl, 100 µM TCEP, 0.001% LMNG).

### Light illumination

Light stimuli were generated with pE-4000 CoolLED for 1 s at the indicated wavelength, using intensity regulators and neutral density filters (ND1-4, Thorlabs, Inc.) to achieve the desired photon flux. Because illumination duration was held constant at 1 s throughout this study, light dose is reported as total photon exposure (photons cm^-2^), calculated as photon flux (photons cm^-2^ s^-1^) × 1 s illumination time; total photon exposures ranged from 10^11^ to 10^16.5^ photons cm^-2^ and are indicated in the figure legends. Photon flux was independently measured using a SpectroCAL MKII spectroradiometer (Cambridge Research Systems) or a S130VC slim photodiode power sensor (UV-extended Si, 200-1100 nm, 500 pW-50 mW; Thorlabs, Inc.) and converted to photon flux and total photon exposure using the measured wavelength and illumination duration. Alternatively, cells were illuminated using the OptoWell system (Opto Biolabs) for 30 s at the following wavelengths and irradiance: 368 nm (0.49 mW cm^-2^), 461 nm (2.74 mW cm^-2^), 519 nm (1.0 mW cm^-2^), corresponding to total photon exposures of approximately 10^16^-10^17^ photons cm^-2^.

### BRET G protein activation assay

HEK293T cells were plated in a 24-well plate (2.5 x10^5^ cells per well, 24 h before transfection) and transiently transfected with plasmid expression vectors for the relevant opsin (250 ng), Gα subunit (250 ng), Gβ-split Venus and Gγ-split Venus subunits (50 ng each), and GRK3-nLuc (12.5 ng)^78^. Cells were detached 6 h after transfection and suspended into 0.5 mL culture medium supplemented with 2 µM retinal and distributed into a 96-well plate (100 µL per well) and incubated at 37°C 5% CO_2_ overnight. Two hours prior to luminescence measurement, the medium was replaced with L-15 medium without phenol red (Thermo Fisher Scientific), supplemented with 1% FBS and 2 µM retinal and cells were kept at room temperature. Ten minutes before measurement, NanoGlo live cell substrate (Promega) diluted in PBS was added (1:400). Wells were measured with an Optima FLUOStar plate reader (BMG) at room temperature and stimulated with light using a custom internal flash setup with the CoolLED (1 s, 10^16^ photons cm^-2^). The BRET ratio was calculated between the measurements collected at 530 nm/450 nm. The data were normalised by dividing everything by the pre-stimulus average and the control conditions (everything except for the receptor DNA).

### GloSensor cAMP assay

HEK293T cells were seeded at density of 2.5 x10^5^ cells per well in 24-well plate 24 h before transfection. Cells were transiently co-transfected with expression vectors of the relevant opsin and pGloSensor-22F cAMP plasmid^80^ at the ratio of 1:1, 500 ng of total DNA per well. Cells from each well were detached with versene 6 h after transfection and suspended into 0.5 mL medium supplemented with an additional 2 µM retinal and distributed into a 96-well plate (100 µL per well) and incubated at 37°C overnight. One hour before measurement, the medium was replaced by assay buffer (phenol-red free L-15 media containing L-glutamine (Gibco), 1% FBS, 2 µM retinal, and 2 mM D-luciferin (MedChemExpress)) and stored at room temperature in the dark. In some cases, no retinal was added to the assay buffer, as specified in the figure legends. Cells were then transferred to the plate reader (PHERAStar, BMG) kept at room temperature, and 1 µM forskolin (Sigma-Aldrich) was added to increase cAMP levels in the chimera screen (Fig. 2d, S4, S5) to see potential Gi inhibitory cAMP responses. After recording a suitable baseline, cells were removed from the plate reader for light or ligand stimulation and were then returned to the reader to record subsequent changes in luminescence.

### Gsx assay

For the Gsx assay, the procedure is the same as the GloSensor cAMP assay, except that HEK293S ΔGs cells were used and additionally 5 ng of individual Gsx G protein chimera DNA was added during transfection^51^. During retinal addition also 125 ng/ml pertussis toxin (Sigma-Aldrich) was added. Response curves were normalised by subtracting the unstimulated condition, then the average of five values after stimulation reached maximum was calculated. These values for each receptor were normalised from 0 to 1.

### Indirect arrestin recruitment assay

Arrestin recruitment was measured with a split-luciferase assay system, in which the large fragment (11s) is anchored to the plasma membrane via a CAAX motif and arrestin is tagged with the small fragment (114)^77^. HEK293T cells were transfected in a 24-well plate 24 h before measurement with 500 ng of total DNA per well with three plasmids in a 1:1:1 ratio (11s-CAAX, 114-arrestin and the receptor). Cells from each well were detached with versene 6 h after transfection and suspended into 0.5 mL medium supplemented with an additional 2 µM retinal and distributed into a 96-well plate (100 µL per well) and incubated at 37°C overnight. One hour before measurement, the medium was replaced by assay buffer (L-15 media (without phenol-red) containing L-glutamine (Gibco), 1% FBS, 2 µM retinal and the cells were stored at room temperature in the dark. Ten minutes before measurement, NanoGlo live cell substrate (Promega) diluted in PBS was added (1:200). Wells were measured with a PHERAStar plate reader (BMG) at room temperature and stimulated with light using the CoolLED (1 s, 10^16^ photons cm^-2^) or by adding 10 µM agonist. To calculate fold change, luminescence values were normalised to the mean of the unstimulated control wells.

### Aequorin calcium assay

HEK293T cells were seeded at a density of 5 x 10^5^ cells per well in a 12-well plate 24 h before transfection. Cells were transiently transfected with vectors of the relevant opsin and the mtAeq at a ratio of 1:1, 500 ng of DNA per well^79^). Cells from each well were detached 6 h after transfection and suspended in 1 mL medium supplemented with an additional 2 µM 11-*cis*-retinal and distributed into a 96-well plate (100 µL per well) and incubated at 37°C with 5% CO_2_ overnight. Two hours prior to luminescence measurement, medium was replaced with L-15 medium without phenol red (Thermo Fisher Scientific), supplemented with 1% FBS, 2 µM 11-*cis*-retinal, and 10 µM Coelenterazine-H (AG Scientific). Cells were incubated and measured at room temperature. Aequorin bioluminescence was measured using an Optima FLUOStar plate reader (BMG) for a total of two minutes. Following 10s baseline measurement, cells were stimulated with light (via CoolLED, 1 s pulse, total photon exposure 10^11.5^ to 10^16^ photons cm^-2^) at six different wavelengths using an integrated internal light stimulus.

### Calculation of opsin photon sensitivity peaks

A dark reference was measured, and other wells were flashed at total photon exposures of 10^11.5^ to 10^16.5^ photons cm^-2^ at six different wavelengths for a duration of 1 s. For JSR1-S199F the Aequorin calcium assay was used, while for optoDRD1 the GloSensor cAMP assay was used. To measure the active-state deactivation spectrum, we pre-activated cells with saturating 385 nm light (10^16^ photons cm^-2^), and allowed cAMP levels to plateau, then measured responses to varying wavelengths. The data was normalised to the pre-stimulus baseline and the maximum response for each light stimulus was extracted and normalised from 0 to 1. The λ_max_ values of opsins were determined using a nonlinear optimization method that minimised the residual error by iteratively fitting the response to the effective photon flux values recalculated with the Govardovskii template nomogram as described previousely^49^.

### Glucose-stimulated insulin secretion GSIS assay

EndoC-βH5® cells (Human Cell Design) were plated in a 96-well plate (1 x10^5^ cells per well) according to the manufacturer’s instructions and cultured in the same well for four weeks at 37°C and 5% CO_2_. Six days after thawing cells were transfected with 50 ng of DNA (EV or optoDRD1) with 0.16 µl Lipofectamine 2000 (Invitrogen). Four hours after transfection the medium was exchanged to starvation medium for 24 h supplemented with 2 µM retinal. The next day GSIS was performed by first exchanging the medium to the βKREBS buffer without glucose and 2 µM retinal and after 1 h the cells were stimulated with 11 mM glucose and additional 10 µM FSK or light stimulation at 385nm or 525 nm for 1 s (10^16^ photons cm^-2^). After 40 minutes at 37°C the medium was collected, and the cells were cultured in the recommended growth medium. One week later, the cells in the same wells were transfected again with the same construct (EV or optoDRD1) and the procedure was repeated. To measure the insulin content the collected medium after stimulation was diluted 50 to 250 times in heat inactivated FBS (tested to have no insulin) and measured with a human insulin ELISA kit (Crystal Chem) according to the manufacturer’s instructions. Values were calculated based on the standard curve and multiplied by the dilution factor.

### Statistical analysis

Data was analysed and plotted using GraphPadPrism 8 software. Data are represented as means ± SEM. For fitting normalised time-resolved signalling assays, either plateau followed by one phase association or decay is used with the least squares regression method. For dose-response experiments, data are normalised and analysed using nonlinear curve fitting for the log (agonist or inhibitor) vs. response (three parameters) curves. Adjusted *p*-values were determined using ordinary one-way ANOVA with Tukey test for multiple comparison (\*\*\*\**p* < 0.0001, \*\*\**p* < 0.001, \*\**p* < 0.01, \**p* < 0.05 were considered statistically significant; *p* > 0.05 was considered statistically not significant (ns)).

### Statistical analysis time-resolved cAMP data

Comparison of parameters extracted from time-resolved cAMP data were performed in Python (version 3.9.13) using pandas, NumPy, SciPy, statsmodels and pingouin; figures were generated with matplotlib. All measurements were treated as independent observations. For each parameter, normality of residuals was assessed with the Shapiro-Wilk test and homogeneity of variances with Levene’s test. If variances were homogeneous (Levene’s test, p ≥ 0.05), groups were compared by one-way ANOVA (ordinary least squares model, type II sums of squares), followed by Tukey’s HSD post hoc test. If variances were unequal (Levene’s test, p < 0.05), Welch’s ANOVA followed by Games-Howell post hoc tests was used instead. The omnibus p-values (ANOVA or Welch’s ANOVA) were additionally adjusted across parameters using the Benjamini-Hochberg procedure; adjusted p < 0.05 was considered statistically significant. Post hoc p-values are corrected for multiple comparisons within each parameter by the respective test. The summary data can be found on Zenodo, parameters_stats.xlsx.

### Matched analysis of chimeras c1-c5

Chimeras c1-c5 were measured in parallel on four independent experimental dates. These data were therefore analysed as a matched (blocked) design, with experimental date as block and n = 4. For each parameter, one-way repeated-measures ANOVA with Greenhouse-Geisser correction was used, and an additive two-way model (y ∼ condition + date) was fitted to quantify the contribution of the experimental date (batch effect). Pairwise comparisons used two-sided paired t-tests with Holm correction within each parameter. The Friedman test was used as non-parametric confirmation, and omnibus *p*-values were adjusted across parameters by Benjamini-Hochberg. Adjusted *p* < 0.05 was considered significant. The summary data can be found on Zenodo, parameters_paired_c1-c5_stats.xlsx.

### AlphaFold2 and 3 predictions

Receptor-miniG protein complex models were predicted with AlphaFold 3 using the AlphaFold Server (https://alphafoldserver.com/). Each prediction contained one receptor chain (chain A) and one mini-G protein chain (chain B). The input sequences are listed in the supplementary (amino acid sequences). Ligands, ions and lipids were not included. Each complex was predicted once with 5 seeds, and all five models were used for analysis. Structures were visualised in PyMOL 2.5.5.

### Extraction of confidence metrics

Confidence scores were extracted from the summary confidence and full data JSON files, and CIF files supplied with each AF3 prediction. Interface residues were defined as receptor-G protein residue pairs with a contact probability above 0.2. For each complex we calculated (i) the number of contacting residue pairs, (ii) the number of unique receptor and unique G protein residues in contact, (iii) the mean contact probability of these pairs, (iv) the mean inter-chain PAE of the same pairs, and (v) the mean pLDDT of the interface residues. Global predicted template modeling score (pTM) and interface pTM (ipTM) were taken directly from the summary file. Analyses were done with a custom Python script available as a Jupyter notebook on Zenodo. pLDDT reports local confidence per residue (0-100). PAE gives the expected position error (in Å) of residue *x* when the predicted structure is aligned on residue *y*. ipTM estimates the accuracy of the predicted relative placement of the two chains and ranges from 0 to 1. The contact probability is the predicted probability that two residues from different chains are in contact, as defined by AF3 (residues in a distance below 8 Å).

### Correlation of structural metrics with signalling

The relationship between AF3 metrics and measured cAMP signalling was tested with Spearman rank correlation (scipy.stats.spearmanr), separately for the initial receptor chimeras and for the c1-c5 series, and for 9-*cis*- and 11-*cis*-retinal conditions. Due to the small number of constructs, we performed leave-one-out analysis and exclusion of selected constructs and corrected for multiple testing across metrics with the Benjamini-Hochberg procedure where indicated. The contact probability threshold determines how many receptor-G protein residue pairs are included in the interface analysis. We tested cut-offs between 0.1 and 0.5 (the resulting contact lists are available as Excel files on Zenodo). Lower thresholds include more residue pairs, which increases the variability between models, whereas higher thresholds exclude genuine interface contacts. We therefore chose 0.2 as a compromise that captures the interface without adding peripheral residues. The overall ranking of the chimeras was unchanged across all tested thresholds.

## Acknowledgements

**Acknowledgments:** We acknowledge Prof. Dr. Raimund Dutzler for co-supervision of the PhD thesis of D.W. and the Biomolecular Structure and Mechanism PhD Program of the Life Science Zurich Graduate School. We thank the National Eye Institute of the National Institutes of Health (NIH) for providing 11-*cis*-retinal. We are grateful to Prof. Dr. Asuka Inoue for kindly providing the HEK293S ΔGs cell line. The plasmids for the arrestin recruitment assay were kindly provided by Dr. Philipp Berger. We thank Prof. Dr. Akihisa Terakita and Prof. Dr. Mitsumasa Koyanagi on input on the S199F mutation of JSR1. This manuscript was peer-reviewed by Review Commons. **Funding:** This work was supported by the European Research Council (ERC) under the European Union’s Horizon 2020 research and innovation program (“ERC SOL,” synergy grant agreement No. 951644 to G.F.X.S.) and the GRC Travel Grant from University of Zurich 2024_Q1_TG_077 to D.W.. **Author Contributions:** D.W. and G.F.X.S designed the project. D.W., R.M., A.P., and H.R., performed experiments. D.W., R.M., A.P. analysed the data. D.W., X.D., R.M, P.I., and G.F.X.S. wrote the paper with input from all authors. **Competing interests:** G.F.X.S. declares that he is a co-founder and scientific advisor of the companies leadXpro AG and InterAx Biotech AG. The other authors declare that they have no competing interests. **Data and material availability:** All data generated or analysed during this study are included in this published article and its supplementary information files. Python analysis scripts, input files and results files are available on Zenodo (DOI: 10.5281/zenodo.24013508). All data shown in the figures are available in Summary_Data.xlsx.

## Supplementary Materials list

Amino acid sequences

Figures S1 to S15

Tables S1 to S6

Summary_Data.xlsx

Zenodo (DOI: 10.5281/zenodo.24013508)

