## Supplementary_data for "Photoswitchable optoGPCRs for reversible control of Gs signalling and arrestin recruitment"

### Amino acid sequences

Following are the coding amino acid sequences of the receptors used in this study including any tags.

> JSR1 B1B1U5, BAG14330.1 kumopsin1 [*Hasarius adansoni*] with N-terminal tag (HA-FLAG-SNAP-TEV-linker-). In bold are ICL2, ICL3 and CT. Yellow highlight shows the S199 position.

MALPVTALLLPLALLLHAARPAAASGIYPYDVPDYAGAGADYKDDDDKGSGSGSGSMDKDCEMKRTTLDSPLGKLELSGCEQGL
HEIKLLGKGTSAADAVEVPAPAAVLGGPEPLMQATAWLNAYFHQPEAIEEFPVPALHHPVFQQESFTRQVLWKLLKVVKFGEVIS
YQQLAALAGNPAATAAVKTALSGNPVPILIPCHRVVSSSGAVGGYEGGLAVKEWLLAHEGHRLGKPGLGPAGSAGENLYFQGGS
GSAGSGSAGSGSAMLPHAAKMAARVAGDHDGRNISIVDLLPEDMLPMIHEHWYKFPPMETSMHYILGMLIIVIGIISVSGNGV
VMYLMMTVKNLRTPGNFLVLNLALSDFGMLFFMMPTMSINCFAETWVIGPFMCELYGMIGSLFGSASIWSLVMITLDRY**NVI
VKGMAGKPL**TKVGALLRMLFVWIWSLGWTIAPMYGWSRYVPEGSMTSCTIDYIDTAINPMSYLIAYAIFVYFVPLFIIIYCYAFI**V
MQVAAHEKSLREQAKKMNIKSLRSNEDNKKASAEFRLAKVA**FMTICCWFMAWTPYLTLSFLGIFSDRTWLTPMTSVWGAIFA
KASACYNPIVY**GISHPKYRAALHDKFPCLKCGSDSPKGDSASTVAESEKAGE**

> JSR1- S199F B1B1U5, BAG14330.1 kumopsin1 [*Hasarius adansoni*] with N-terminal tag (HA-FLAG-SNAP-TEV-linker-). In bold are ICL2, ICL3 and CT. Yellow highlight shows the F199 position.

MALPVTALLLPLALLLHAARPAAASGIYPYDVPDYAGAGADYKDDDDKGSGSGSGSMDKDCEMKRTTLDSPLGKLELSGCEQGL
HEIKLLGKGTSAADAVEVPAPAAVLGGPEPLMQATAWLNAYFHQPEAIEEFPVPALHHPVFQQESFTRQVLWKLLKVVKFGEVIS
YQQLAALAGNPAATAAVKTALSGNPVPILIPCHRVVSSSGAVGGYEGGLAVKEWLLAHEGHRLGKPGLGPAGSAGENLYFQGGS
GSAGSGSAGSGSAMLPHAAKMAARVAGDHDGRNISIVDLLPEDMLPMIHEHWYKFPPMETSMHYILGMLIIVIGIISVSGNGV
VMYLMMTVKNLRTPGNFLVLNLALSDFGMLFFMMPTMSINCFAETWVIGPFMCELYGMIGSLFGSASIWSLVMITLDRY**NVI
VKGMAGKPL**TKVGALLRMLFVWIWSLGWTIAPMYGWSRYVPEGSMTFCTIDYIDTAINPMSYLIAYAIFVYFVPLFIIIYCYAFI**V
MQVAAHEKSLREQAKKMNIKSLRSNEDNKKASAEFRLAKVA**FMTICCWFMAWTPYLTLSFLGIFSDRTWLTPMTSVWGAIFA
KASACYNPIVY**GISHPKYRAALHDKFPCLKCGSDSPKGDSASTVAESEKAGE**

ICL2, ICL3, ICL4-helix8, CT after palmitoylation site. These parts have been used to design the DRD1 chimera with varying amount of loops exchanged. 1D4 tag

> JSR1_DRD1 based on: P21728|DRD1_HUMAN D(1A) dopamine receptor
MLPHAAKMAARVAGDHDGRNISIVDLLPEDMLPMIHEHWYKFPPMETSMHYILGMLIIVIGIISVSGNGVVMYLMMTVKNL
RTPGNFLVLNLALSDFGMLFFMMPTMSINCFAETWVIGPFMCELYGMIGSLFGSASIWSLVMITLDRYWAISSPFRYERKMTKV
GALLRMLFVWIWSLGWTIAPMYGWSRYVPEGSMTSCTIDYIDTAINPMSYLIAYAIFVYFVPLFIIIYCYAFIYRIAQKQIRRIAALER
AAVHAKNCQTTTGNGKPVECSQPESSFKMSFKRETKVLKTLFMTICCWFMAWTPYLTLSFLGIFSDRTWLTPMTSVWGAIFAK
ASACYNPIVYAFNADFRKAFSTLLGCYRLCPATNNAIETVSINNNGAAMFSSHHEPRGSISKECNLVYLIPHAVGSSEDLKKEEAAG
IARPLEKLSPALSVILDYDTDVSLEKIQPITQNGQHPTGSTETSQVAPA

> JSR1_5HT7R based on: P34969|5HT7R_HUMAN 5-hydroxytryptamine receptor 7
MLPHAAKMAARVAGDHDGRNISIVDLLPEDMLPMIHEHWYKFPPMETSMHYILGMLIIVIGIISVSGNGVVMYLMMTVKNLR
TPGNFLVLNLALSDFGMLFFMMPTMSINCFAETWVIGPFMCELYGMIGSLFGSASIWSLVMITLDRYLGITRPLTYPVRQTKVGA
LLRMLFVWIWSLGWTIAPMYGWSRYVPEGSMTSCTIDYIDTAINPMSYLIAYAIFVYFVPLFIIIYCYAQIYKAARKSAAKHKFPGF
PRVEPDSVIALNGIVKLQKEVEECANLSRLLKHERKNISIFKREQKAATTLFMTICCWFMAWTPYLTLSFLGIFSDRTWLTPMTSV
WGAIFAKASACYNPIVYGIFNRDLRTTYRSLLQCQYRNINRKLSAAGMHEALKLAERPERPEFVLQNADYCRKKGHDSGSTETSQ
VAPA

> JSR1_AA2BR based on: P29275|AA2BR_HUMAN Adenosine receptor A2b
MLPHAAKMAARVAGDHDGRNISIVDLLPEDMLPMIHEHWYKFPPMETSMHYILGMLIIVIGIISVSGNGVVMYLMMTVKNLR
TPGNFLVLNLALSDFGMLFFMMPTMSINCFAETWVIGPFMCELYGMIGSLFGSASIWSLVMITLDRYLAICVPLRYKSLVTKVGA

LLRMLFVWIWSLGWTIAPMYGWSRYVPEGSMTSCTIDYIDTAINPMSYLIAYAIFVYFVPLFIIIYCYAFIFLVACRQLQRTELMDHS
RTTLQREIHAAKSLAMTICCWFMAWTPYLTLSFLGIFSDRTWLTPMTSVWGAIFAKASACYNPIVYAYRNRDFRYTFHKIISRYLLC
QADVKSGNGQAGVQPALGVGLGSTETSQVAPA

> JSR1_ β2AR based on: P07550|ADRB2_HUMAN Beta-2 adrenergic receptor
MLPHAAKMAARVAGDHDGRNISIVDLLPEDMLPMIHEHWYKFPPMETSMHYILGMLIIVIGIISVSGNGVVMYLMMTVKNLR
TPGNFLVLNLALSDFGMLFFMMPTMSINCFAETWVIGPFMCELYGMIGSLFGSASIWSLVMITLDRYNAITSPFKYQSLLTKVGA
LLRMLFVWIWSLGWTIAPMYGWSRYVPEGSMTSCTIDYIDTAINPMSYLIAYAIFVYFVPLFIIIYCYARVFQEAKRQLQKIDKSEG
RFHVQNLSQVEQDGRTGHGLRRSSKFCLKEHKALKTLFMTICCWFMAWTPYLTLSFLGIFSDRTWLTPMTSVWGAIFAKASACY
NPIVYGIRSPDFRIAFQELLCLRRSSLKAYGNGYSSNGNTGEQSGYHVEQEKENKLLCEDLPGTEDFVGHQGTVPSDNIDSQGRN
CSTNDSLLGSTETSQVAPA

> JSR_LSHR based on: P22888 · LSHR_HUMAN Lutropin-choriogonadotropic hormone receptor
MLPHAAKMAARVAGDHDGRNISIVDLLPEDMLPMIHEHWYKFPPMETSMHYILGMLIIVIGIISVSGNGVVMYLMMTVKNLR
TPGNFLVLNLALSDFGMLFFMMPTMSINCFAETWVIGPFMCELYGMIGSLFGSASIWSLVMITLERWHTITYAIHLDQKLRKVG
ALLRMLFVWIWSLGWTIAPMYGWSRYVPEGSMTSCTIDYIDTAINPMSYLIAYAIFVYFVPLFIIIYCYAKIYFAVRNPELMATNKD
TKIAKKMFMTICCWFMAWTPYLTLSFLGIFSDRTWLTPMTSVWGAIFAKASACYNPIVYAIFTKTFQRDFFLLLSKFGCCKRRAEL
YRRKDFSAYTSNCKNGFTGSNKPSQSTLKLSTLHCQGTALLDKTRYTECGSTETSQVAPA

> JSR1_JellyOp_1 ICL3 only based on: B6F0Y5_CARRA *Carybdea rastonii* (Box jellyfish)
MLPHAAKMAARVAGDHDGRNISIVDLLPEDMLPMIHEHWYKFPPMETSMHYILGMLIIVIGIISVSGNGVVMYLMMTVKNLR
TPGNFLVLNLALSDFGMLFFMMPTMSINCFAETWVIGPFMCELYGMIGSLFGSASIWSLVMITLDRYNVIVKGMAGKPLTKVG
ALLRMLFVWIWSLGWTIAPMYGWSRYVPEGSMTSCTIDYIDTAINPMSYLIAYAIFVYFVPLFIIIYCYALVQGEMKNMRGRAAQ
LFGSESEAALKNIKAEKRHTKVAFMTICCWFMAWTPYLTLSFLGIFSDRTWLTPMTSVWGAIFAKASACYNPIVYGISHPKYRAAL
HDKFPCLKCGSDSPKGDSASTVAESEKAGEGSTETSQVAPA

> JSR1_Taar9 based on: Q96RI9|TAAR9_HUMAN Trace amine-associated receptor 9
MLPHAAKMAARVAGDHDGRNISIVDLLPEDMLPMIHEHWYKFPPMETSMHYILGMLIIVIGIISVSGNGVVMYLMMTVKNLR
TPGNFLVLNLALSDFGMLFFMMPTMSINCFAETWVIGPFMCELYGMIGSLFGSASIWSLVMITLDRYIAVTDPLTYPTKFTKVGA
LLRMLFVWIWSLGWTIAPMYGWSRYVPEGSMTSCTIDYIDTAINPMSYLIAYAIFVYFVPLFIIIYCYAKIFLVAKHQARKIESTASQ
AQSSSESYKERVAKRERKAAKTLGIAICCWFMAWTPYLTLSFLGIFSDRTWLTPMTSVWGAIFAKASACYNPIVYAFFYQWFGKA
IKLIVSGKVLRTDSSTTNLFSEEVETDGSTETSQVAPA

**Sequences used only for AlphFold predictions**

>Mini-Gs_393_Nehme2017

GIEKQLQKDKQVYRATHRLLLLGADNSGKSTIVKQMRILHGGSGGSGGTSGIFETKFQVDKVNFHMFDVGGQRDERRKWIQC
FNDVTAIIFVVDSSDYNRLQEALNDFKSIWNNRWLRTISVILFLNKQDLLAEKVLAGKSKIEDYFPEFARYTTPEDATPEPGEDPR
VTRAKYFIRDEFLRISTASGDGRHYCYPHFTCAVDTENARRIFNDCRDIIQRMHLRQYELL

> mini-Giq04 mini-G protein based on the jumping spider sequence

MGHHHHHHENLYFQMIDRNLREDGEKARREVKLLLLGADNSGKSTIVKQMKIIHGGSGGSGGTTGIVETHFTFKSIHFKMFDV
GGQRSERKKWIHCFEDVTAIIFCVDLSDYNRMHESMKDFDSICNNKWFTDTSIILFLNKKDLFEEKIKKSPLTICYPEYAGSNTYEE
AAAYIQCQFEDLNKRKDTKEIYTHFTCATDTKNARLIFC AVKDTILQNNLKECNLV

### Supplementary figures

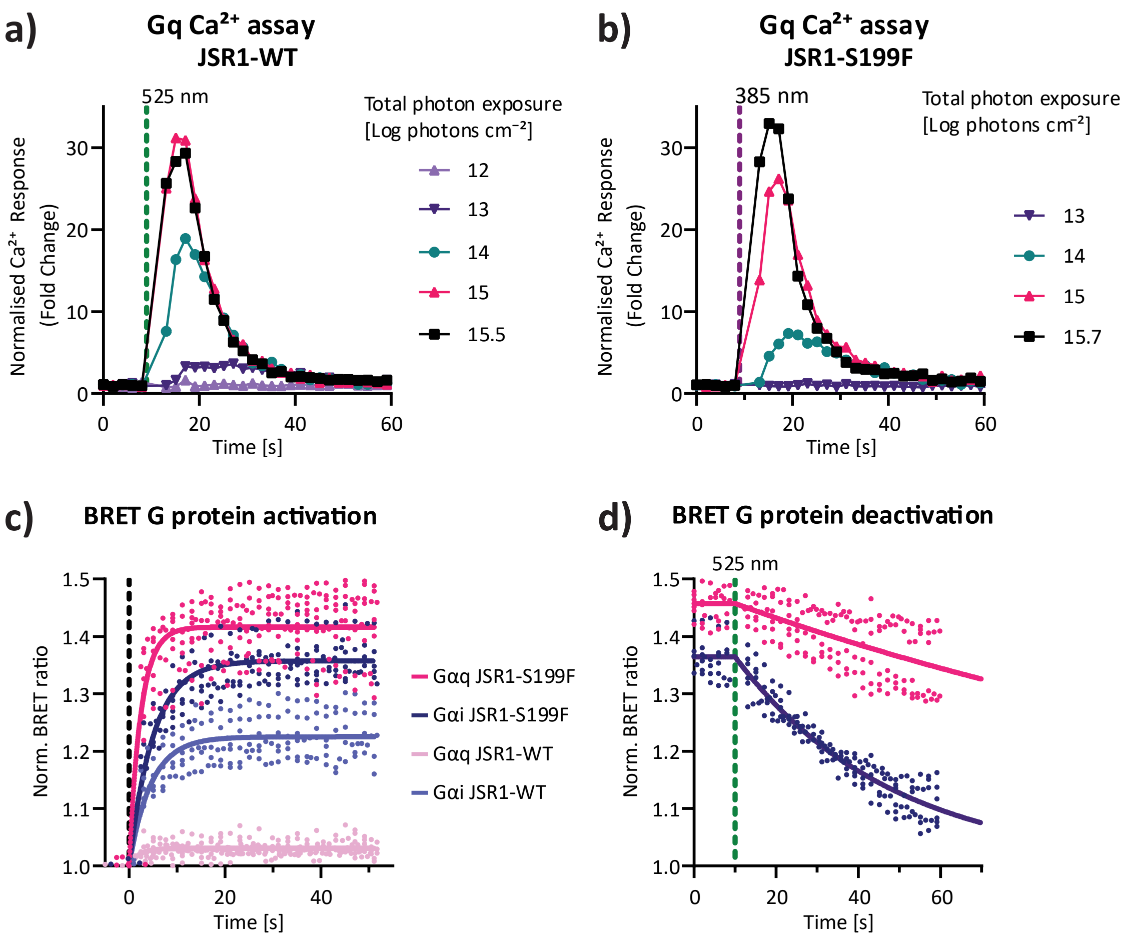

e)

| | | | k [s-1] | SEM |
|---|---|---|---|---|
| **activation** | **JSR1-WT** | **Gq** | 0.42 | 0.36 |
| | | **Gi** | 0.22 | 0.02 |
| | **JSR1-S199F** | **Gq** | 0.41 | 0.06 |
| | | **Gi** | 0.20 | 0.01 |
| **deactivation** | **JSR1-S199F** | **Gq** | 0.006 | 0.003 |
| | | **Gi** | 0.026 | 0.007 |

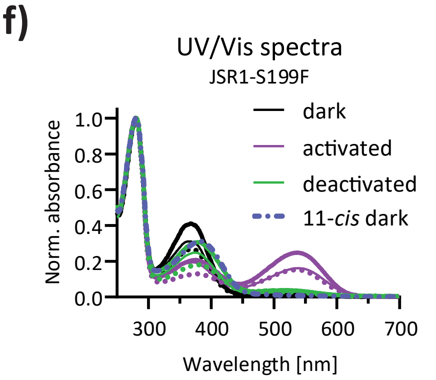

**Figure S1. Gi and Gq signalling profiles of JSR1-WT and the JSR1-S199F mutant in HEK293T cells. a, b)** Gq signalling activity measured as increase in intracellular $Ca^{2+}$ recorded using a bioluminescent aequorin sensor. Both JSR1-WT and JSR1-S199F induce a similar $Ca^{2+}$ response after 1 s of light stimulus (JSR1-WT (525 nm) and JSR1-S199F mutant (385 nm)). **c, d)** Gαi and Gαq activity measured by BRET G protein activation assay and kinetic fits including three biological replicates for Gq data and two biological replicates for Gi data each including three technical replicates. **c)** Activation of JSR1-WT and JSR1-S199F with 525 nm and 385 nm light respectively ($10^{15.5}$ photons $cm^{-2}$). **d)** Deactivation of JSR1-S199F after 525 nm light stimulation (green dashed line; $10^{15.5}$ photons $cm^{-2}$). **e)** Activation and deactivation rates obtained from fits to the curves in panels c and d. The rate of activation of Gq is about two times faster than for Gi, while the deactivation is about two times slower for Gq than for Gi. The data correspond to the mean and standard error (SEM) of individual fits of 6-9 replicates (2-3 biological replicates). **f)** UV-vis spectra of JSR1-S199F of n = 3 independent protein purifications with 9-*cis*-retinal, or 11-*cis*-retinal when indicated. The colour indicates if it was measured before light exposure (dark) or after activation with violet light or deactivation with green light, and the line texture indicates replicates (rep. 1: 0.5 pt line, rep. 2: dotted line, rep. 3: 2 pt line).

**a)**

| GPCRdb position | 2x39 | 3x49 | 3x50 | 3x53 | 3x54 | 3x55 | 34x50 | 34x51 | 34x53 | 34x54 | 34x55 | 5x61 | 5x64 | 5x65 | 5x68 |
|---|---|---|---|---|---|---|---|---|---|---|---|---|---|---|---|
| Sequence Consensus | T | D | R | A | I | T | | | | | | V | E | A | Q |
| Sequence Conservation | 63 | 81 | 99 | 80 | 80 | 45 | 55 | 55 | 55 | 55 | 56 | 41 | 41 | 67 | 60 |
| Property Consensus | Hb | +- | +- | SN | HA | Hb | | | | | | HA | Hb | HA | Hb |
| Property Conservation | 67 | 100 | 100 | 88 | 99 | 63 | 55 | 55 | 55 | 55 | 56 | 92 | 61 | 95 | 77 |
| Interaction Conservation | 41 | 24 | 97 | 95 | 97 | 49 | 45 | 45 | 29 | 35 | 23 | 72 | 49 | 88 | 70 |

| GPCRdb position | 5x69 | 5x71 | 5x72 | 5x75 | 5x76 | 6x29 | 6x32 | 6x33 | 6x36 | 6x37 | 7x56 | 8x47 | 8x48 | 8x49 |
|---|---|---|---|---|---|---|---|---|---|---|---|---|---|---|
| Sequence Consensus | L | K | I | | | | K | A | T | L | | S | P | D |
| Sequence Conservation | 38 | 38 | 61 | 38 | 45 | 61 | 77 | 50 | 54 | 80 | 61 | 45 | 37 | 47 |
| Property Consensus | HA | Hb | HY | Hb | Hb | | Hb | HY | Hb | HY | | Hb | HY | Hb |
| Property Conservation | 69 | 67 | 65 | 42 | 52 | 61 | 94 | 77 | 76 | 96 | 61 | 86 | 48 | 81 |
| Interaction Conservation | 48 | 41 | 70 | 28 | 38 | 23 | 61 | 94 | 82 | 69 | 23 | 33 | 23 | 21 |

**b)**

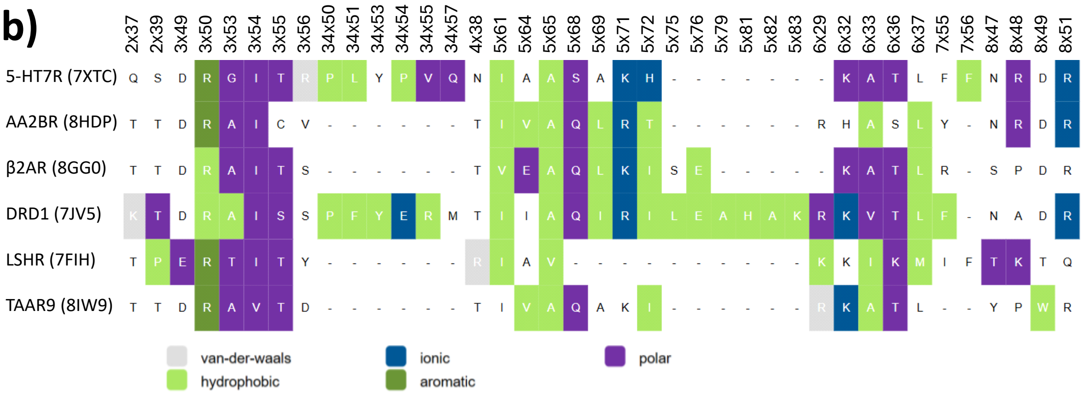

| GPCRdb position | 2x37 | 2x39 | 3x49 | 3x50 | 3x53 | 3x54 | 3x55 | 3x56 |
|---|---|---|---|---|---|---|---|---|
| Sequence Consensus | T | T | D | R | A | I | T | S |
| Sequence Conservation | 67 | 67 | 83 | 100 | 67 | 83 | 67 | 33 |
| Property Consensus | Hb | Hb | +- | +- | Sm | HA | Hb | Hb |
| Property Conservation | 100 | 83 | 100 | 100 | 83 | 100 | 83 | 83 |
| Interaction Conservation | 17 | 33 | 17 | 100 | 100 | 100 | 83 | 17 |

| GPCRdb position | 4x38 | 5x61 | 5x64 | 5x65 | 5x68 | 5x69 | 5x71 | 5x72 |
|---|---|---|---|---|---|---|---|---|
| Sequence Consensus | T | I | | A | Q | | K | I |
| Sequence Conservation | 67 | 83 | 33 | 83 | 67 | 33 | 50 | 50 |
| Property Consensus | Hb | HA | HA | HA | Hb | HA | +- | HA |
| Property Conservation | 100 | 100 | 83 | 100 | 83 | 83 | 83 | 50 |
| Interaction Conservation | 17 | 83 | 50 | 100 | 83 | 50 | 67 | 83 |

| GPCRdb position | 6x29 | 6x32 | 6x33 | 6x36 | 6x37 | 7x55 | 7x56 | 8x47 | 8x48 | 8x49 | 8x51 |
|---|---|---|---|---|---|---|---|---|---|---|---|
| Sequence Consensus | R | K | A | T | L | F | | N | | D | R |
| Sequence Conservation | 50 | 83 | 67 | 67 | 83 | 33 | 67 | 50 | 33 | 67 | 83 |
| Property Consensus | +- | +- | HA | Hb | HA | HY | | Hb | αH | Hb | Hb |
| Property Conservation | 67 | 100 | 100 | 100 | 100 | 67 | 67 | 100 | 67 | 83 | 100 |
| Interaction Conservation | 50 | 67 | 100 | 83 | 67 | 17 | 17 | 17 | 50 | 17 | 50 |

**c)**

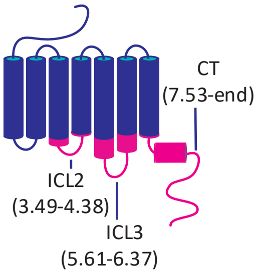

**d)**

ICL2

| GPCRdb no. | 3.48 | **3.49** | **3.50** | **3.51** | 3.52 | 3.53 | 3.54 | 34.54 | 34.55 | 34.56 | 34.57 | 4.38 | 4.39 | 4.40 |
|---|---|---|---|---|---|---|---|---|---|---|---|---|---|---|
| JSR1 | L | **D** | **R** | **Y** | N | V | I | K | P | L | _ | T | K | V |
| JSR1-DRD1 | L | **D** | **R** | **Y** | W | A | I | E | R | K | M | T | K | V |
| DRD1 | V | **D** | **R** | **Y** | W | A | I | E | R | K | M | T | P | K |

ICL3

| GPCRdb no. | 5.58 | 5.59 | 5.60 | 5.61 | 5.62 | 5.63 | 5.64 | 6.35 | 6.36 | 6.37 | 6.38 | 6.39 | 6.40 | 6.41 |
|---|---|---|---|---|---|---|---|---|---|---|---|---|---|---|
| JSR1 | Y | A | F | I | V | M | Q | K | V | A | F | M | T | I |
| JSR1-DRD1 | Y | A | F | I | Y | R | I | K | T | L | F | M | T | I |
| DRD1 | Y | T | R | I | Y | R | I | K | T | L | S | V | I | M |

CT

| GPCRdb no. | **7.49** | **7.50** | 7.51 | 7.52 | **7.53** | 7.54 | 7.55 | 7.56 | 8.47 | 8.48 |
|---|---|---|---|---|---|---|---|---|---|---|
| JSR1 | **N** | **P** | I | V | **Y** | G | I | S | H | P |
| JSR1-DRD1 | **N** | **P** | I | V | **Y** | A | F | _ | N | A |
| DRD1-WT | **N** | **P** | I | I | **Y** | A | F | _ | N | A |

**Figure S2. Consensus GPCR-Gs protein interface map. a)** The table, obtained from the GPCRdb (34,35) shows the GPCR residues forming contacts with the Gs protein in at least 20% of the structures. Data include experimental structures of human Class A Gs-coupled receptors without fusions (n = 186). **b)** All GPCR residues forming contacts with the Gs protein extracted from GPCRdb from the cryo-EM structures of the Gs-coupled receptors included in this study: β2AR (PDB ID: 8GG0), DRD1 (PDB ID: 7JV5), AA2BR (PDB ID: 8HDP), 5-HT7R (PDB ID: 7XTC), TAAR9 (PDB ID: 8IW9), and LSHR (PDB ID: 7FIH). In the upper panel the sequences are displayed, and the background colour of the amino acids specifies the type of interaction it has to the Gs protein as specified in the legend. The bottom table displays the consensus sequence of those six Gs receptors and to which extend the amino acid properties are conserved. The codes in 'Property Consensus' are: hydrophobic (HY), hydrophobic aliphatic (HA), charged (+-), small (Sm), α-helix propensity (αH), and hydrogen bonding ability (Hb). **c)** Scheme of the JSR1 chimeras in which ICL2, ICL3, ICL4, helix 8, and the C-terminus (CT) are exchanged with the sequence of a Gs-receptor (pink) to redirect its native signalling properties towards Gs signalling. For simplicity we call the exchanged regions ICL2, ICL3 and CT although they also contain part of the ordered alpha-helical domains, as apparent from their GPCRdb number 3.49 (helix 3 position 49), 34.55 (loop between helix 3 and 4). **d)** Sequence alignment of the exchanged regions between JSR1, DRD1, and the JSR1-DRD1 chimera including all identified contact sites in ICL2, ICL3 and ICL4. The two contact sites in ICL1 (2.37 and 2.39) were not included.

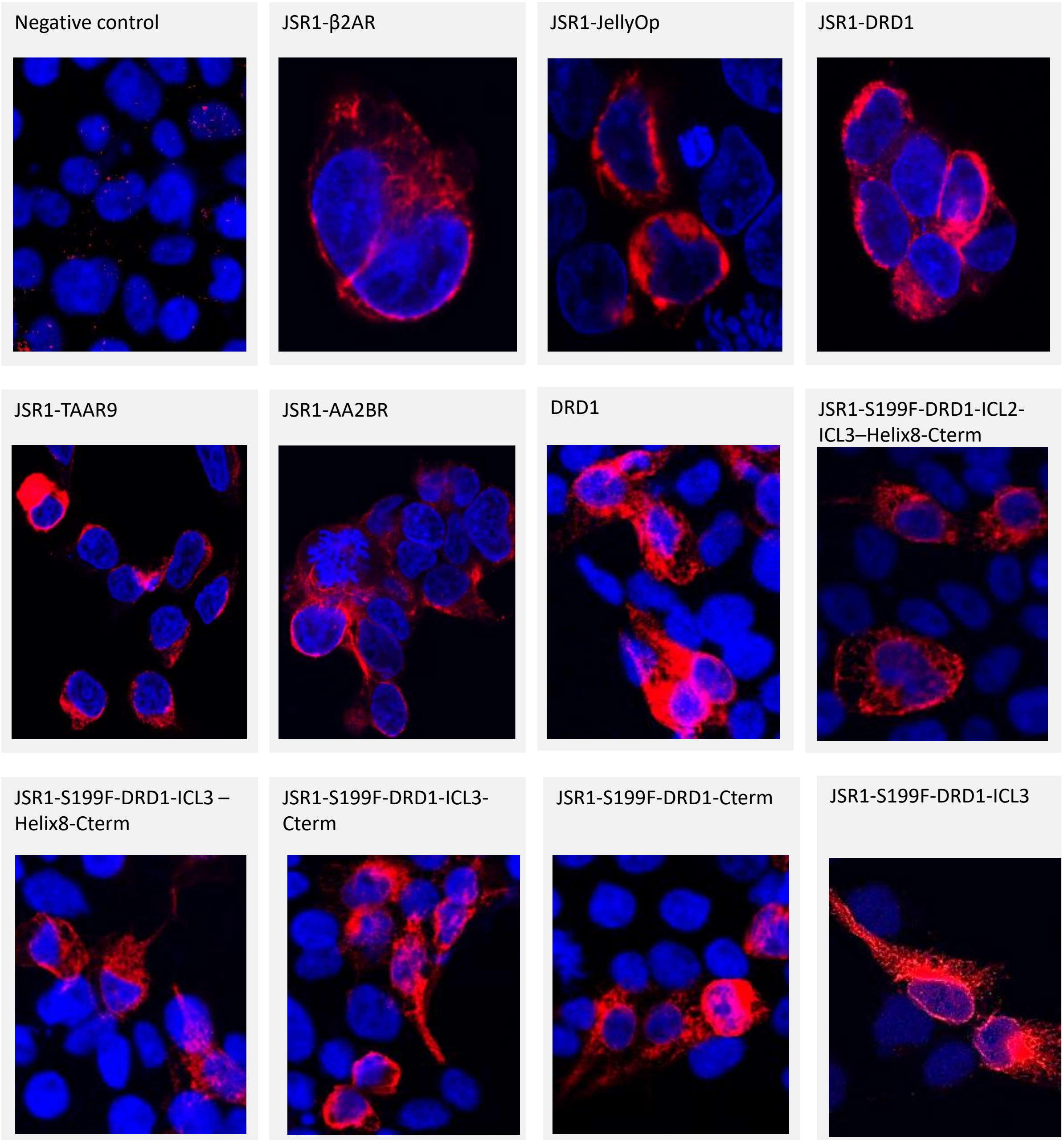

**Figure S3. Immunocytochemistry assays of chimeric receptors.** All the chimeric receptor constructs have a Rho-1D4 tag at the C-terminus. HEK293T cells were fixed 24h after transfection and labelled with a 1D4 antibody and a secondary antibody labelled with Alexa Fluor 594 (red). The nucleus was stained with DAPI (blue).

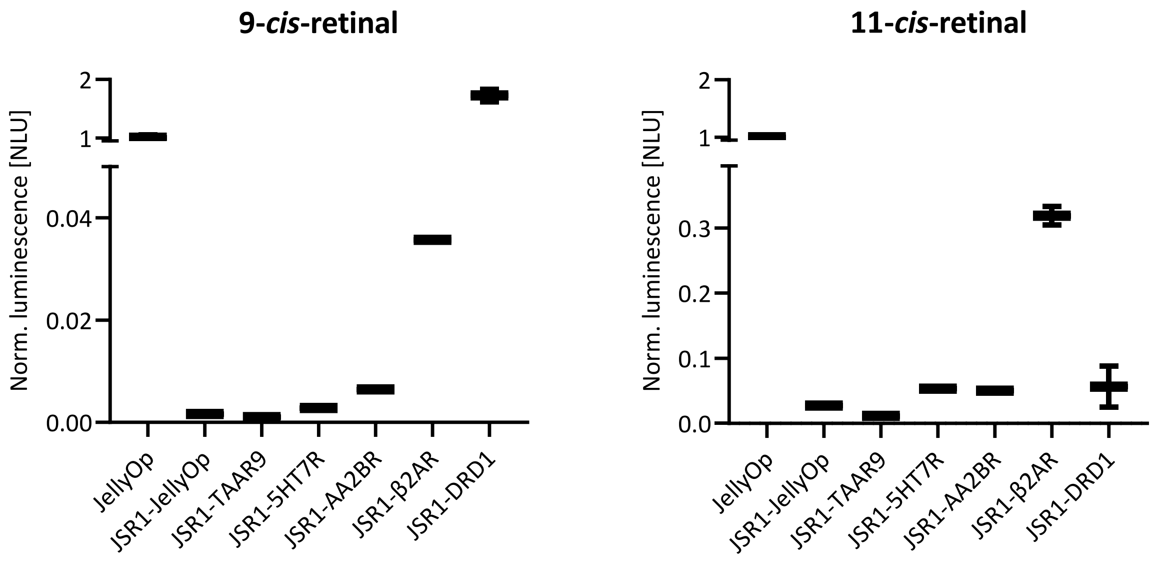

**Figure S4. Response maxima of the GloSensor cAMP assay in HEK293T cells.** Time-resolved curves are depicted in Figure S5. After subtracting the unstimulated baseline, we plotted the maximum response to light stimulation (OptoWell, 519 nm, $10^{17}$ photons $cm^{-2}$) in the cAMP assay for each chimera supplemented with 9-*cis*- and 11-*cis*-retinal. All 9-*cis*-retinal conditions (except for JellyOp) were measured with 1 µM forskolin, therefore preventing direct comparison to JellyOp in this figure. Direct comparison between optoDRD1 and JellyOp show that they reach similar amplitudes in the cAMP assay (Fig. 4f, Fig. S14, and Fig. 5a). The JSR1-DRD1 chimera produces the strongest response with 9-*cis*-retinal; however, with 11-*cis*-retinal it shows a high baseline activity with no further increase upon light stimulation (Fig. S5). A similar retinal-specific effect is also observed in the JSR1-S199F-DRD1 chimera (Fig. S12), although in this case the high basal activity can be switched off with green light. Most of the other chimeras have a higher signal amplitude when supplemented with 11-*cis*-retinal, leading to the hypothesis that the intracellular domains allosterically modulate the retinal binding site.

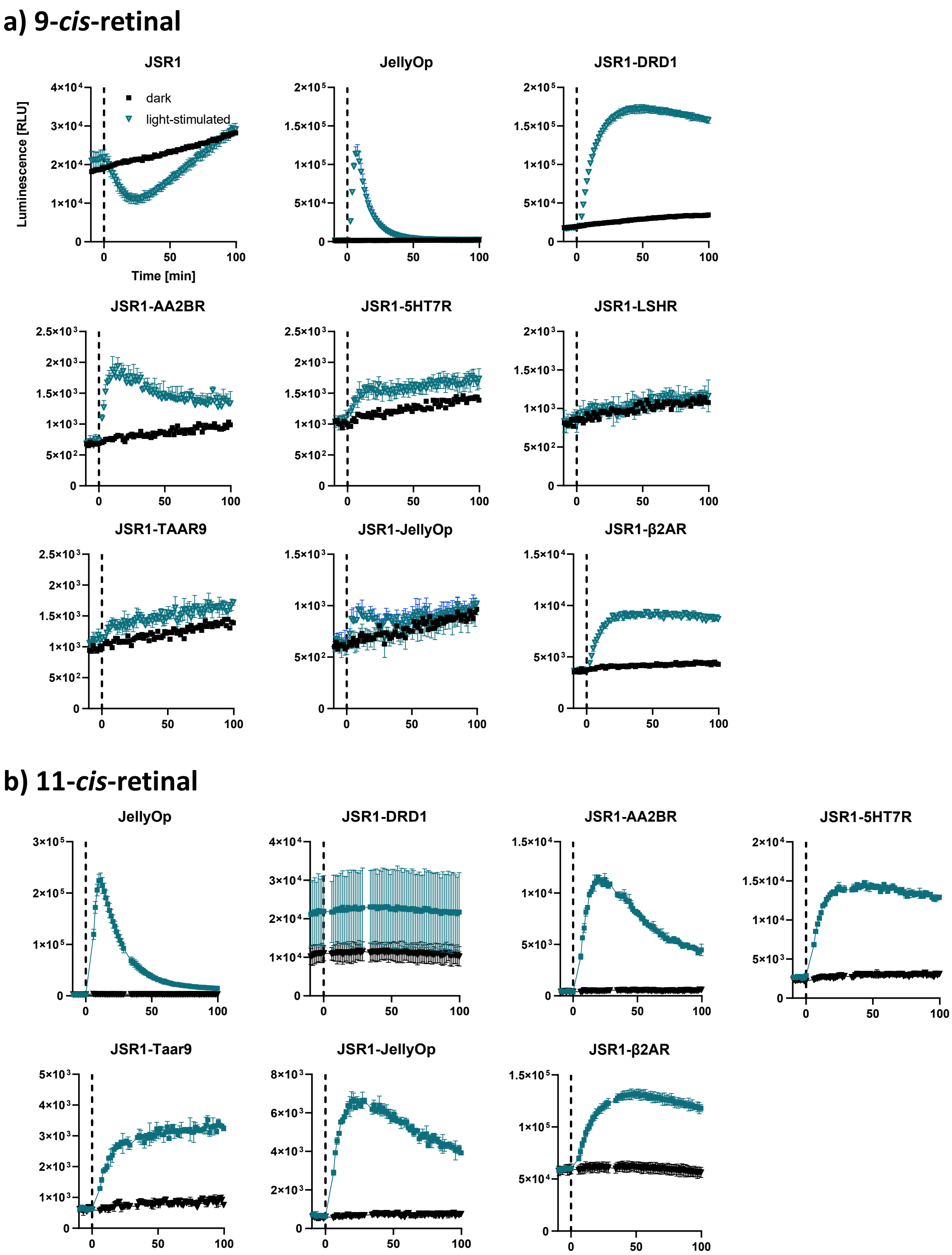

**Figure S5. GloSensor cAMP assay in HEK293T cells for JSR1 chimeras and controls (JSR1-WT, JellyOp).** Light stimulation was applied at T = 0. **a)** Cells were supplemented with 9-*cis*-retinal. Forskolin (1 µM) was added to all 9-*cis*-retinal conditions (except for JellyOp) to allow comparison with JSR1-WT. **b)** Cells were supplemented with 11-*cis*-retinal. The kinetic profiles vary: some chimeras sustain prolonged cAMP signals, while others decay rapidly. Decay of activity may be linked to desensitization through arrestin.

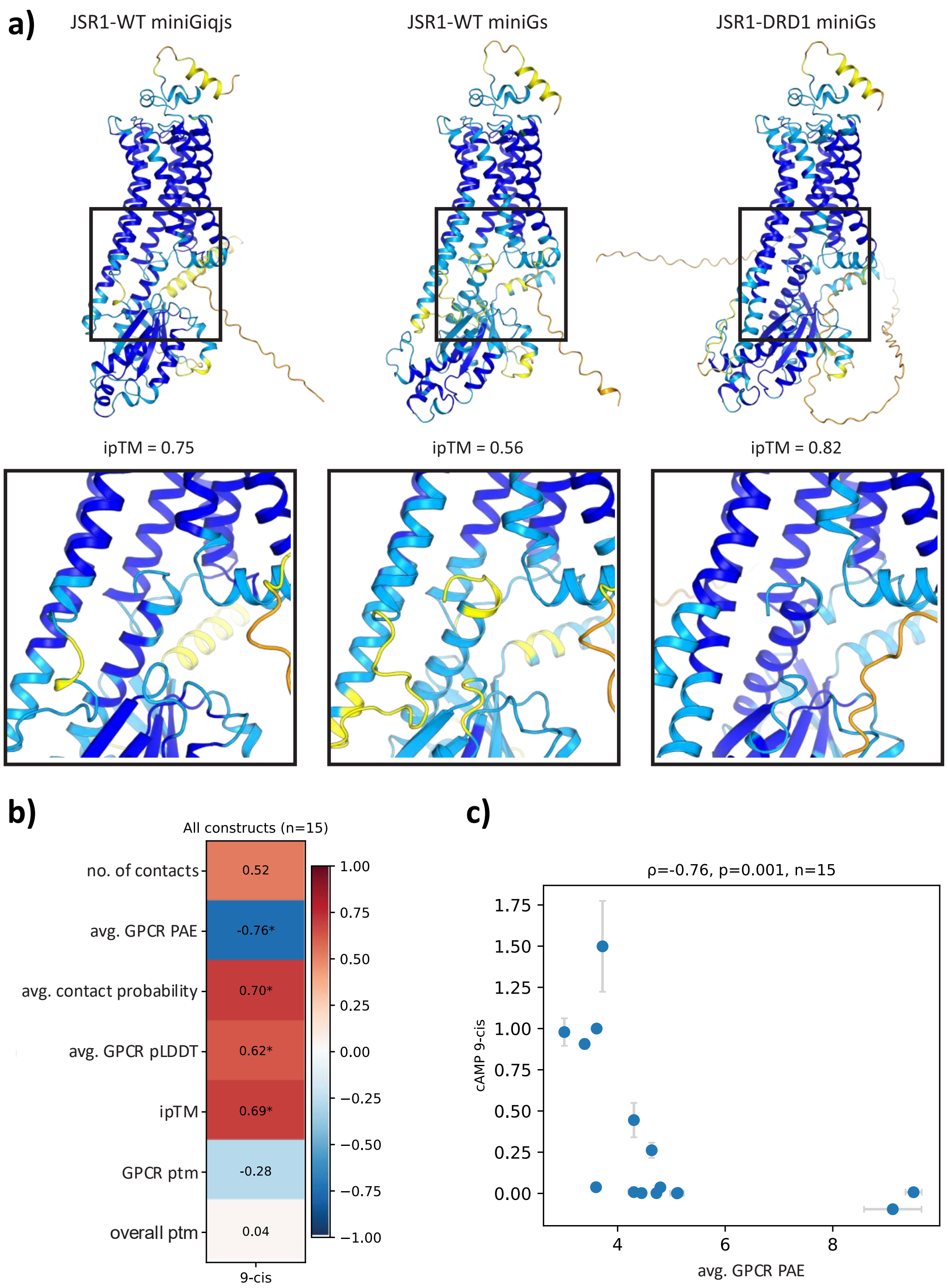

**Figure S6. a)** Comparison of AF models of different GPCR/miniG protein complexes. The models are coloured according to their pLDDT scores (dark blue: > 90%; light blue: 70-90%; yellow: 50-70%, orange: <50%). JSR1-WT does not bind Gs proteins, which agrees with its lower pLDDT score for this G protein binding interface (central panels) compared to the other two models. The lower affinity is also apparent in the lower iPTM score of the JSR1-WT miniGs model. **b)** AlphaFold interface-confidence metrics for all constructs modelled with miniG proteins to the maximum amplitude of cAMP signalling data measured with GloSensor for cells supplemented with 9-*cis*-retinal (JSR1, initial chimera screening constructs, c1-c5, DRD1 and JellyOp). The metrics are defined in Methods (l. 145–184). **c)** Scatter plot of the AF-fold parameter with the highest correlation to the measured cAMP activity for cells supplemented with 9-*cis*-retinal.

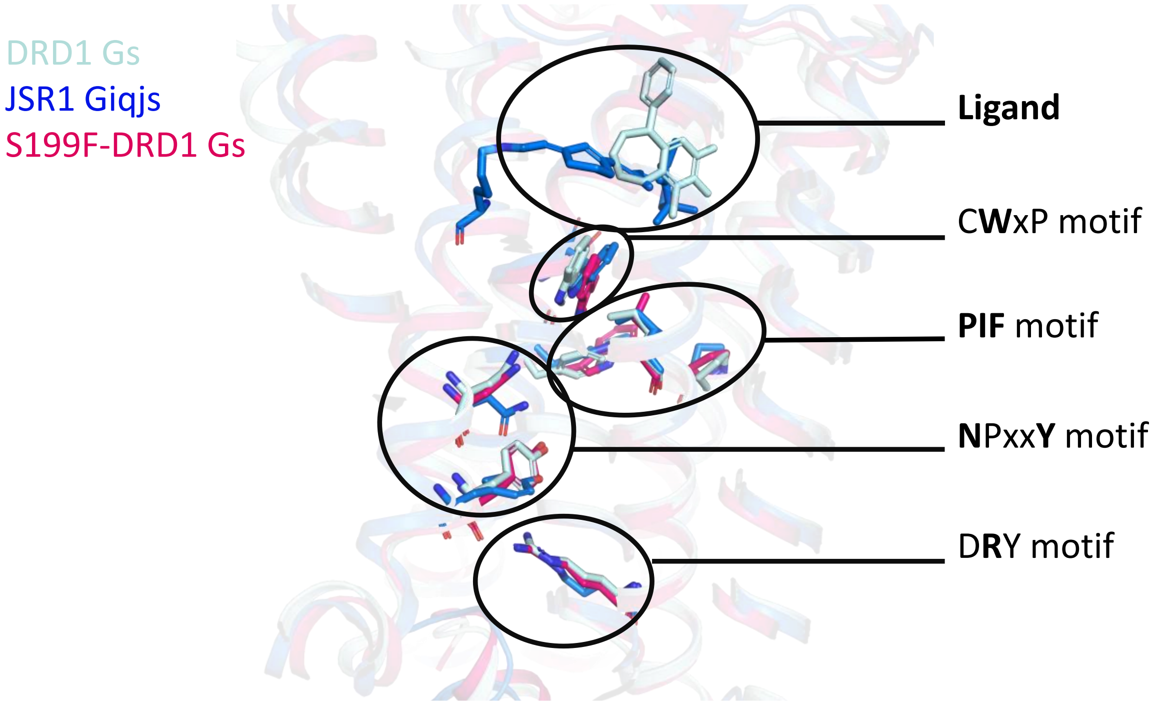

**Figure S7.** Comparison between the active state structure of the JSR1-jumping spider Giq protein complex (PDB ID: 9EPQ, blue), the DRD1-Gs protein complex structure (PDB ID: 7JV5; cyan), and the AF prediction of JSR1-S199F-DRD1 chimera (red). The residues of selected conserved motifs in Class A GPCRs are shown as sticks. The activation mechanism of JSR1 starts with the isomerization of the 11-*cis*-, or 9-*cis*- to all-*trans*-retinal. Subsequently, the conserved $W^{6.48}$ of the CWxP ($C^{6.47}$, $W^{6.48}$, $P^{6.50}$) motif pushes on the $W^{6.44}$, which is part of the PIF ($P^{5.50}$, $I^{3.40}$, $F^{6.44}$) motif. Most of the receptors that were included in the chimera design feature the conserved $F^{6.44}$ at this position, while JSR1 harbours $W^{6.44}$. Finally, the ionic lock between $R^{3.50}$-$E^{6.30}$ breaks. TM6 moves outward and the $R^{3.50}$ of the DRY ($D^{3.49}$, $R^{3.50}$, $Y^{3.51}$) motif establishes a hydrogen bond to the C-terminus of the Gα subunit (Y393 H5.23).

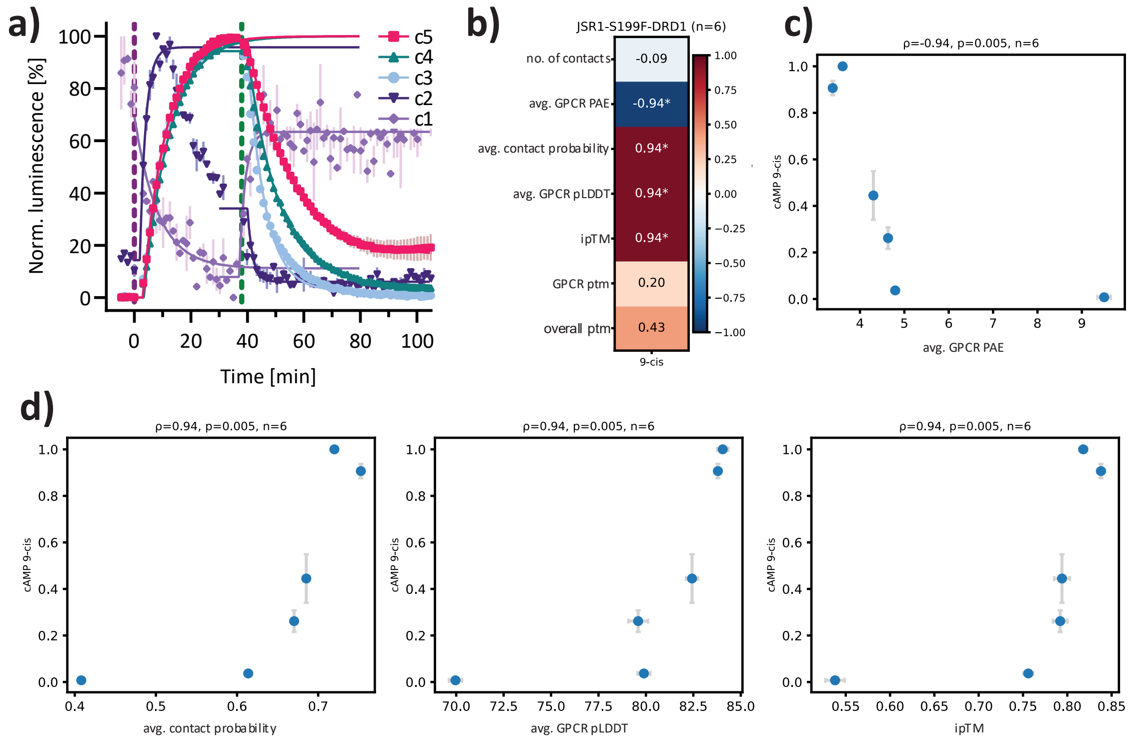

**Figure S8**. GloSensor cAMP responses of JSR1-S199F DRD1 chimeras in HEK293T cells, with fitted activation and deactivation curves. **a)** Normalised time-resolved cAMP responses from Fig. 3b; mean ± SEM, n = 2. The fitted line is shown as a solid line in the same colour. Fitted parameters for all biological replicates are provided in Figure S9. **b)** Correlation matrix of AF parameters to the maximum amplitude of cAMP signalling data for c1-c5 chimeras and DRD1. **c-d)** Scatter plot of the AF-fold parameters with the highest correlation to the measured cAMP activity for c1-c5 chimeras and DRD1. **c)** negative correlation (PAE), **d)** positive correlation (contact probability, pLDDT, ipTM).

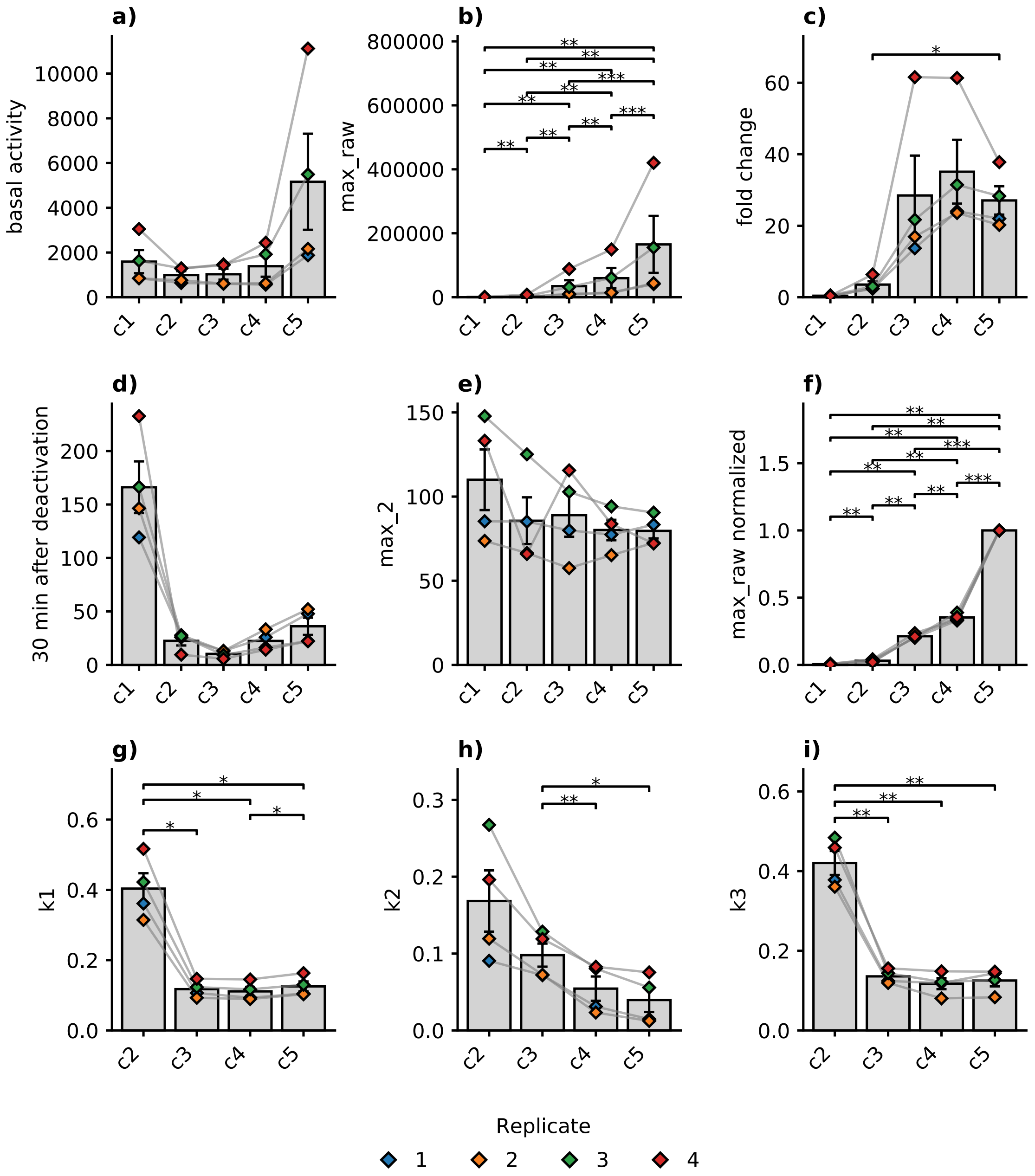

**Figure S9**. GloSensor cAMP responses of JSR1-S199F DRD1 chimeras in HEK293T cells with extracted parameters. Pairwise comparisons were done by two-sided paired t-tests with Holm adjustment, brackets indicate significant pairs (* $p < 0.05$, ** $p < 0.01$, *** $p < 0.001$). Individual values are shown as diamonds and are colored by replicate number; grey lines connect measurements from the same date. **a)** Raw basal activity before stimulation. **b)** Max_raw is the maximum raw value extracted between 0 and 37 minutes. **c)** Fold change of maximal response, calculated as the maximum divided by the pre-stimulation baseline. **d)** Percentage of the value at 30 minutes after deactivation compared to the maximum value. **e)** The second maximum value extracted after T2 (second activation) normalized to the first maximum value. **f)** The maximum raw values normalized to the c5 condition of the same date measured in the same plate. **g-i)** Rate constants that have been fitted to the activation of the receptor (rise of cAMP) and deactivation (fall of cAMP). c1 was excluded from this comparison since it has Gi activity and an opposite effect on cAMP (activation leads to a decrease of cAMP)**. g)** $k_1$ is the rate constant extracted from the rising phase of cAMP. **h)** $k_2$ is the rate constant from the falling phase of cAMP. **i)** $k_3$ is the rate constant for the second rising phase when receptors were reactivated. Fitted parameters are provided in Table S5.

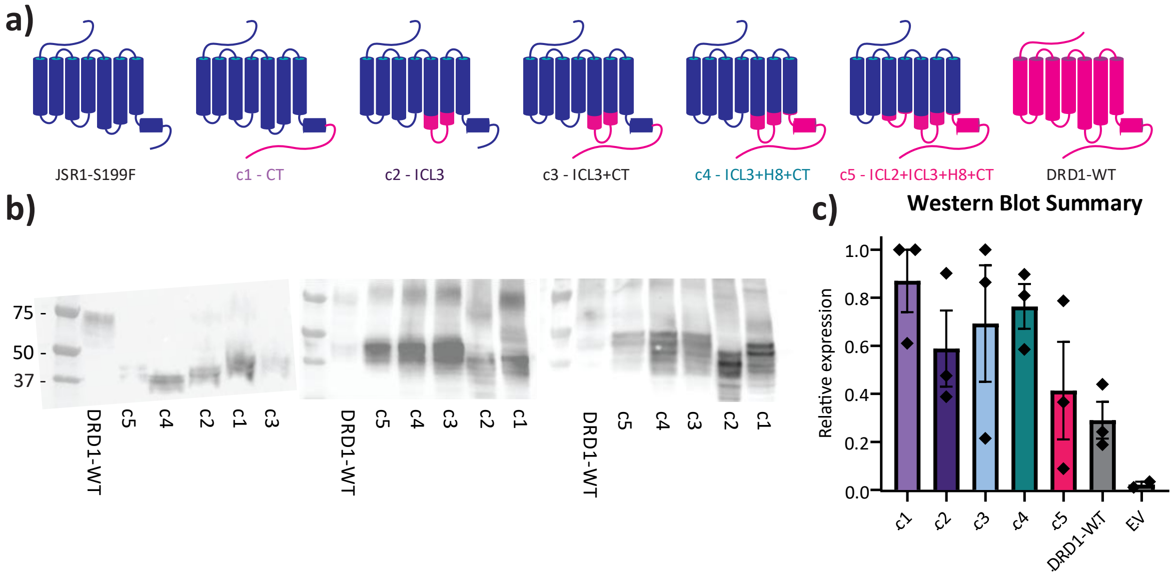

**Figure S10**. Expression levels of different JSR1-S199F-DRD1 chimeras in HEK293T cells. **a)** Cartoon depiction of the engineered JSR1-S199F-DRD1 chimeras (blue: JSR1, pink: DRD1). **b)** Western blot bands of the chimeras from three independent experiments**.** Monomeric receptors are expected to be between 35-50 kDa apparent molecular weight, differences in size are expected due to longer sequence of the ICL3 and the CT of the DRD1. The bands around 75 -100 kDa may represent receptors with posttranslational modifications including glycosylation and lipidation[1] or dimeric species and were also included into the quantification. **c)** Quantification of the intensity of the Western blot bands mean ± SEM, n = 3.

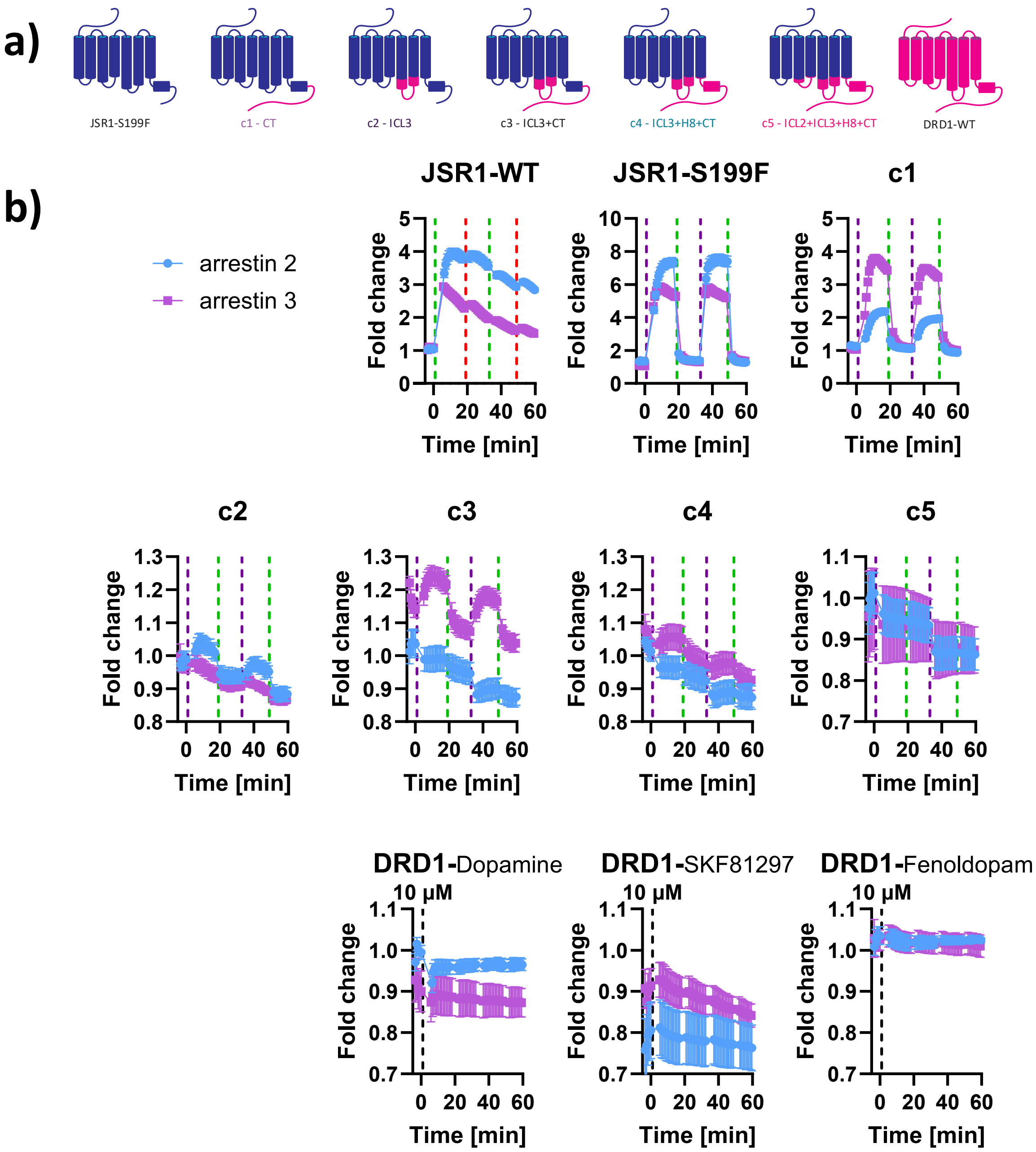

**Figure S11**. Indirect arrestin recruitment assay in HEK293T cells supplemented with 9-*cis*-retinal. **a)** Cartoon depiction of the engineered JSR1-S199F-DRD1 chimeras. The pink segments are from DRD1, and blue are from JSR1-S199F. **b)** Cells were stimulated either once with 10 µM agonist or with successive 1 s light pulses at $10^{16}$ photons $cm^{-2}$ at the time points indicated by dashed lines (green: 525 nm, red: 595 nm, violet: 385 nm). Data are presented as mean ± SEM fold change relative to unstimulated controls (n = 3). JSR1-WT, JSR1-S199F and c1 show robust light-induced arrestin recruitment that is reversed by deactivating illumination in JSR1-S199F and c1. Chimera c2 shows preferential recruitment of arrestin 2 relative to arrestin 3, whereas c3 displays the opposite preference. Notably, arrestin 2 recruitment to JSR1-S199F is more dependent on the C-terminus of JSR1, while arrestin 3 recruitment is mainly dependent on the ICL3 of JSR1. Substituting both ICL3 and the C-terminus to DRD1 leads to some arrestin 3 recruitment.

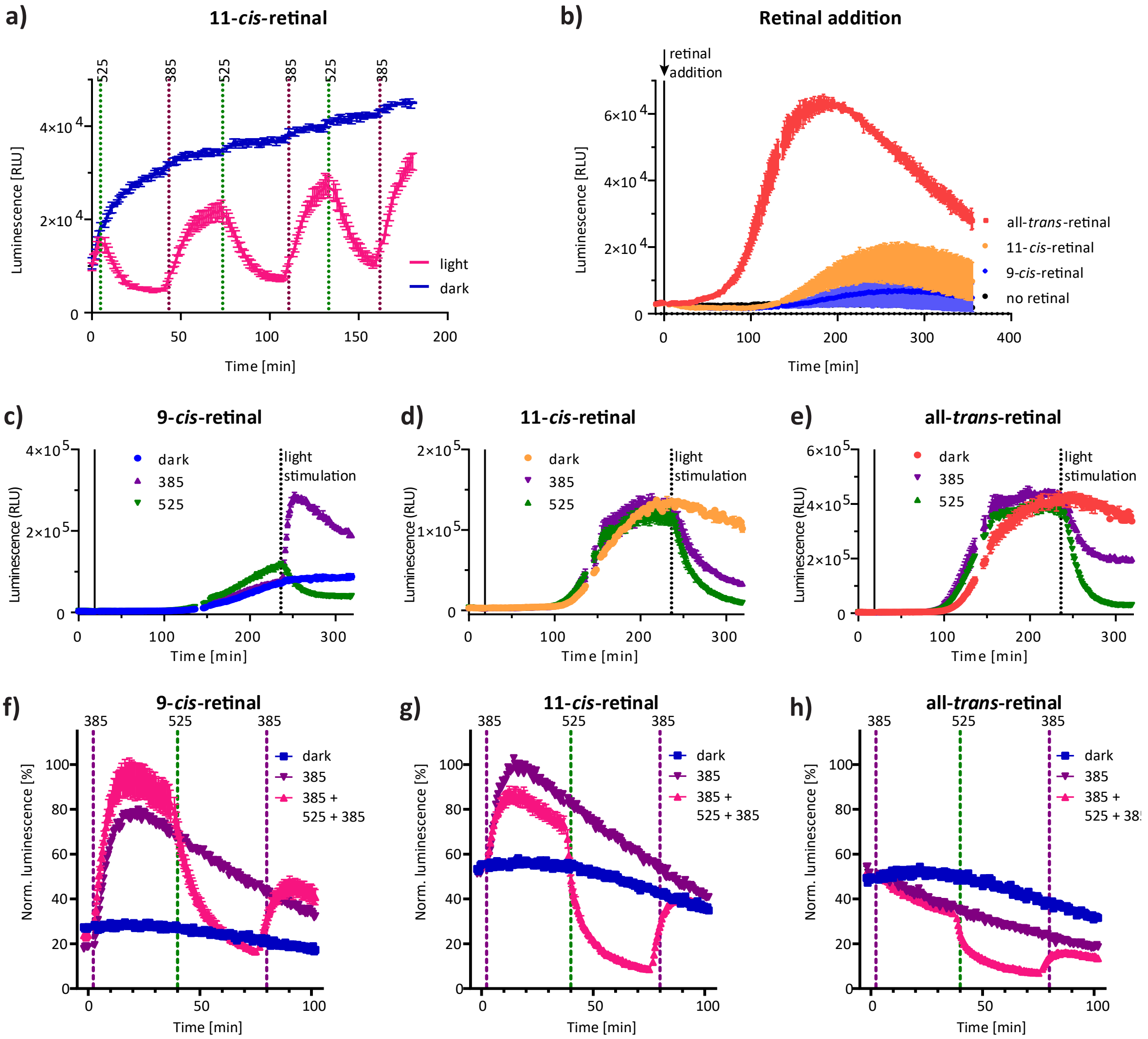

**Figure S12.** Comparison of JSR1-S199F-DRD1 c5 (optoDRD1) supplemented with different retinal chromophores in a GloSensor cAMP assay. **a)** HEK293T cells with optoDRD1 and 11-*cis*-retinal supplemented overnight in the medium and in the assay buffer (10 µM). Wells were stimulated alternating with green light (525 nm) and violet light (385 nm) with $10^{16}$ photons $cm^{-2}$ at the indicated time points. **b)** Different retinal isoforms (9-*cis*-, 11-*cis*-, all-*trans*-retinal) were added at T = 0 (final concentration: 10 µM) to HEK293T cells expressing optoDRD1, n = 2. **c-e)** Retinal isoforms (10 µM) were added to HEK293T cells expressing optoDRD1 only at the beginning of the measurement (solid line), and wells were subsequently kept in the dark or stimulated with green or violet light (dashed line), n = 2. All-*trans*-retinal led to the highest activation over time, followed by 11-cis- and 9-cis-retinal. When adding 11-cis- or all-trans-retinal, stimulation with either green or violet light led to a decrease in the signal, suggesting that only all-*trans*-retinal binds to the active state of the opsin to render it photosensitive. However, when adding 9-*cis*-retinal, the signal increased with violet light and decreased with green light. This suggests that binding of 9-*cis*-retinal yields a majority of receptors that are activatable with violet light and a smaller fraction that is already active and can be deactivated with green light (probably a subpopulation bound to all-*trans*-retinal formed by the isomerization of 9-*cis*-retinal). **f-h)** Neuro-2a cells with optoDRD1. 10µM retinal was added overnight to the medium and in the assay buffer. Cells were stimulated as indicated in the figure legends with $10^{16}$ photons $cm^{-2}$, n = 3.

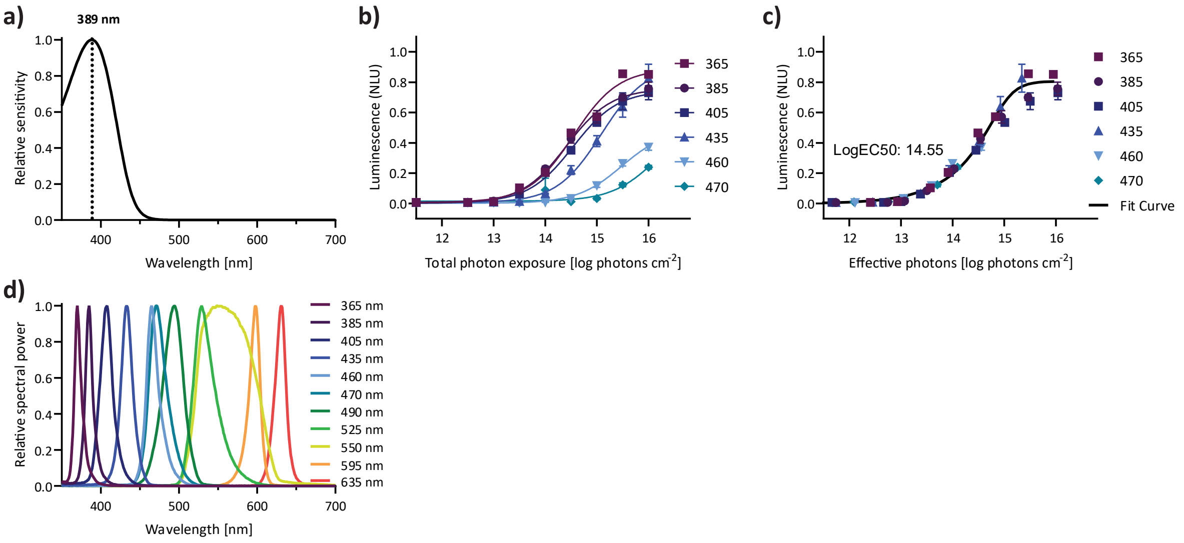

**Figure S13.** Light sensitivity of JSR1-S199F measured in a spectral sensitivity assay in HEK293T cells. **a)** Govardovskii plot of signalling responses across six different wavelengths at ten different intensities each. The resulting $\lambda_{max}$ value is 389 ± 0.54 nm for activation. **b)** Dose-response curves for 1 s receptor illumination across intensities ranging from $10^{11}$ to $10^{16}$ photons cm$^{-2}$ at the specified wavelengths. **c)** Normalised dose response curves to effective photon flux, calculated based on the determined $\lambda_{max}$. Data represent the mean ± SEM of three biological replicates. **d)** Power spectra of the illumination wavelengths from the CoolLED pE4000, which was used to stimulate the receptor and record the action spectra.

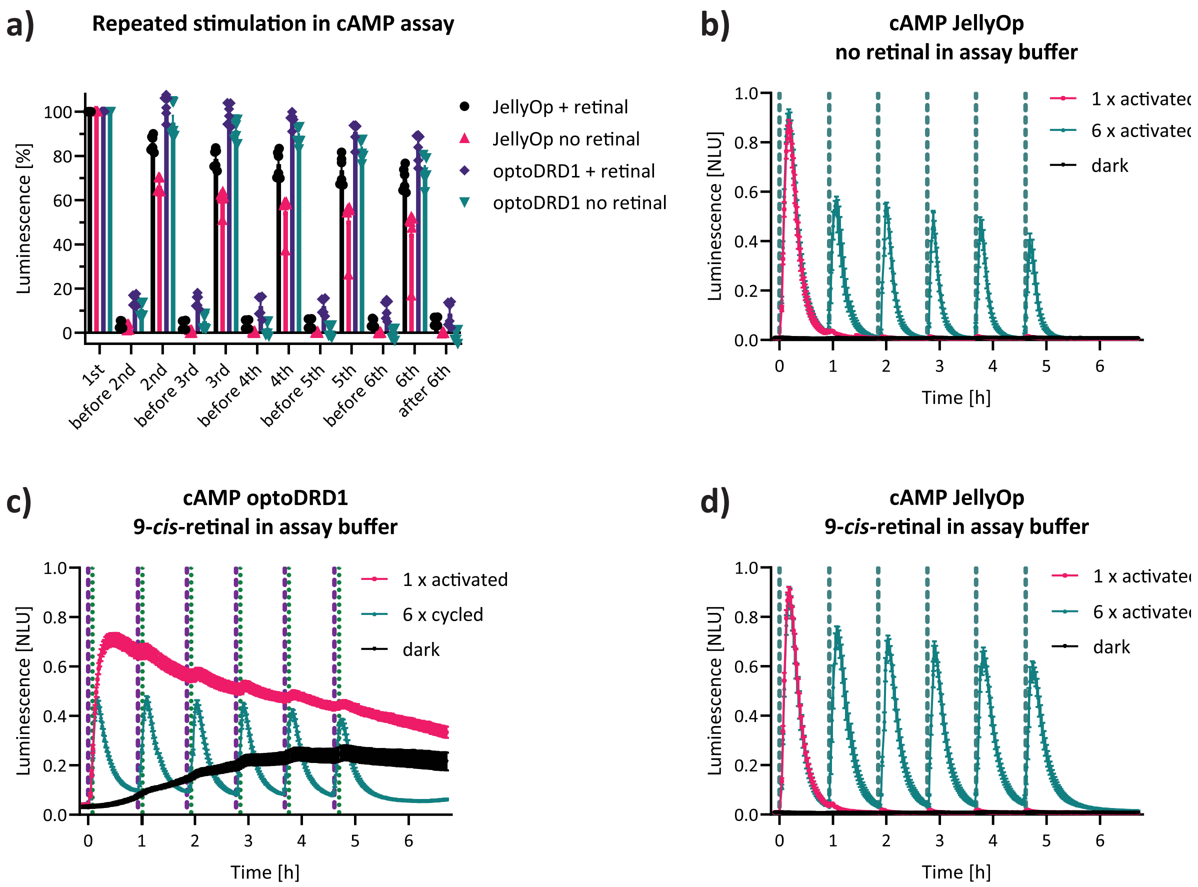

**Figure S14.** Comparison of optoDRD1 and JellyOp in matched experimental conditions. Transfected HEK293T cells were supplemented over night with 2 µM 9-*cis*-retinal and either the assay buffer was supplemented with 2 µM retinal or no retinal was added in the assay buffer (retinal-limited condition). **a)** The average of three values was extracted from the time-resolved curves at the peak and just before the next stimulation (~50 minutes later). Values are plotted as percentages of the first stimulation peak for each receptor and individual replicate ((x/baseline-1)/(1$^{st}$/baseline-1) *100%). Anova of all optoDRD1 peaks show no significant difference between conditions with and without retinal (p-adj. between 0.22 and 0.051), while all peaks of JellyOp without retinal were significantly lower than the conditions with retinal (p-adj. < 0.0005). **b-d)** Values were normalized to the highest luminescence value of each biological replicate (same date and plate), including both the JellyOp and optoDRD1 values of Fig. 4f and S13b-d (the highest value was in one of the JellyOp wells). **b)** JellyOp without additional retinal in the assay buffer. It is repeatably activatable but there is a marked decrease of peak amplitude after the 1$^{st}$ stimulus. **c)** OptoDRD1 with additional 9-*cis*-retinal in the assay buffer. OptoDRD1 was activated with violet light and deactivated 5 minutes later with green light. This pattern was repeated six times. The baseline is increasing during the measurement, and the deactivation is not as complete as in the optoDRD1 conditions without retinal in the assay buffer (**Fig. 4f**). **d)** JellyOp with 9-*cis*-retinal in the assay buffer. The second peak is decreased in intensity, but less than in conditions without retinal addition to the assay buffer.

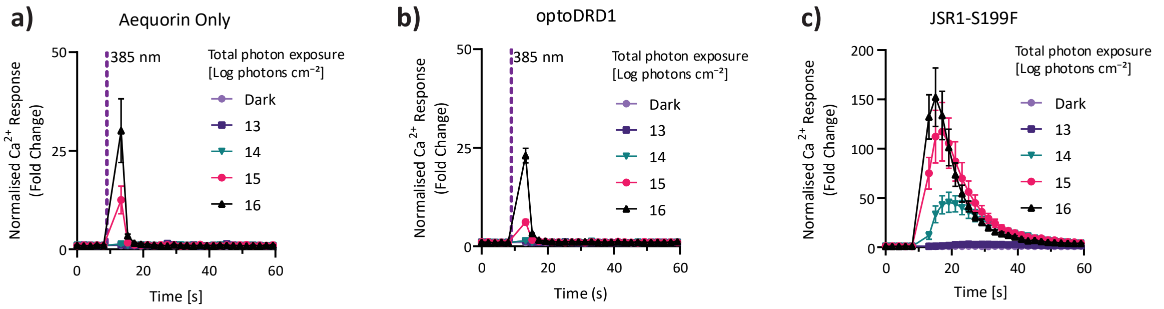

**Figure S15.** Assessment of Gq activity of optoDRD1 with aequorin assay in HEK293T cells. **a)** Cells expressing only the aequorin sensor were stimulated with 385 nm light at different intensities. At high intensities, a transient increase in luminescence was observed in a single measurement point (<10 s), which returned immediately to baseline. This light-induced artifact is distinct from a receptor-dependent $Ca^{2+}$ response, which is defined as luminescence remaining elevated over several consecutive measurement points (~30 s), as seen in c. **b)** Cells expressing optoDRD1 supplemented with 9-*cis*-retinal were stimulated at different light intensities. No Gq activated $Ca^{2+}$ response could be detected. Data represent the mean ± SEM, n=4. **c)** Cells expressing JSR1-S199F supplemented with 9-*cis*-retinal were stimulated at different light intensities. Data represent the mean ± SEM, n=3.

### Supplementary tables

**Table S1.** Length of intracellular segments of the receptors used in this study. Long segments are highlighted in pink and short ones in blue.

| Receptors | ICL2 length (3.52-4.38) | ICL3 length (5.60-6.37) | CT length after NPxxY (7.54-end) |
|---|---|---|---|
| 5HT7R | 14 | 69 | 59 |
| AA2BR | 14 | 33 | 42 |
| β2AR | 14 | 55 | 87 |
| DRD1 | 14 | 59 | 115 |
| JellyOP | 15 | 38 | 45 |
| JSR1 | 13 | 44 | 41 |
| LSHR | 14 | 24 | 76 |
| TAAR9 | 14 | 42 | 37 |

**Table S2. Summary of the seven chimera designs (JSR1 + Gs-coupled receptors).** Start and end positions of the insertions are provided in GPCRdb numbering. In case of matching overlapping residues between JSR1 and the Gs-coupled receptor, the reported stretches correspond to the maximal sequence that is present in the Gs-coupled receptor. The two right columns rate the signal amplitude with 9-*cis*- and 11-*cis*-retinal as displayed in Fig. 2d, S4 and S5.

| | ICL2 | | | ICL3 | | | C-term | | cAMP signal 9-*cis*-retinal | cAMP signal 11-*cis*-retinal |
|---|---|---|---|---|---|---|---|---|---|---|
| loops from | yes/ no | start | end | yes/ no | start | end | yes/ no | start | order of best (1) to worst (6) | order of best (1) to worst (6) |
| DRD1 | yes | 3.49 | 34.56 | yes | 5.61 | 6.37 | yes | 7.53 | 1 | Active before light stimulation |
| 5HT7R | yes | 3.49 | 34.56 | yes | 5.6 | 6.37 | yes | 7.56 | 4 | 2 |
| TAAR9 | yes | 3.49 | 4.38 | yes | 5.6 | 6.4 | yes | 7.53 | 5 | 5 |
| LSHR | yes | 3.49 | 4.38 | yes | 5.6 | 6.37 | yes | 7.53 | No signal | No signal |
| β2AR | yes | 3.53 | 34.56 | yes | 5.6 | 6.37 | yes | 7.56 | 2 | 1 |
| JellyOp | no | NA | NA | yes | 5.6 | 6.34 | no | NA | 6 | 4 |
| AA2BR | yes | 3.49 | 34.56 | yes | 5.61 | 6.39 | yes | 7.49 | 3 | 3 |

**Table S3. Interface confidence parameters of the AlphaFold predictions for the WT Gs coupling receptors and JSR1-Gs chimeras.** The table displays the number of contact points predicted to have a contact probability of > 0.2 and their average (avg.) predicted aligned error (PAE) for the receptor. Additionally, the average contact probability and the chain-interface predicted template modelling values (ipTM) are shown. The data are colour coded from green to red based on their values. For PAE, lower values indicate better predictions and are shown in green (overall range: 3.01–9.7). In contrast, for number of contacts, avg. contact probability, pLDDT and iPTM, higher values represent better predictions and are displayed in green. The values are given as mean and SD of the five models provided by a single AlphaFold prediction with five seeds.

| | Signalling | Constructs | number of contacts > 0.2 | | Avg. Receptor PAE | | Avg. contact probability | | Avg. Receptor pLDDT | | ipTM | |
|---|---|---|---|---|---|---|---|---|---|---|---|---|
| | Signalling | Constructs | M | SD | M | SD | M | SD | M | SD | M | SD |
| WT | Gs | 5HT7R + miniGs | 104 | 0.00 | 5.56 | 0.08 | 0.62 | 0.00 | 75.34 | 0.27 | 0.81 | 0.01 |
| | | AA2BR + miniGs | 74 | 0.00 | 3.62 | 0.05 | 0.65 | 0.00 | 84.70 | 0.26 | 0.81 | 0.00 |
| | | β2AR + miniGs | 80 | 0.00 | 3.69 | 0.10 | 0.71 | 0.00 | 84.66 | 0.56 | 0.79 | 0.01 |
| | | DRD1 + miniGs | 100 | 0.00 | 3.39 | 0.03 | 0.75 | 0.00 | 83.79 | 0.18 | 0.84 | 0.00 |
| | | JellyOP + miniGs | 85 | 0.00 | 3.01 | 0.05 | 0.66 | 0.00 | 89.60 | 0.28 | 0.80 | 0.01 |
| | | LSHR + miniGs | 64 | 0.00 | 4.61 | 0.06 | 0.65 | 0.00 | 78.75 | 0.23 | 0.81 | 0.00 |
| | | TAAR9 + miniGs | 86 | 0.00 | 3.65 | 0.04 | 0.69 | 0.00 | 85.97 | 0.15 | 0.81 | 0.00 |
| chimeras | Gs (expected) | JSR1-5HT7R + miniGs | 90 | 0.00 | 5.12 | 0.05 | 0.66 | 0.00 | 74.84 | 0.30 | 0.77 | 0.00 |
| | | JSR1-AA2BR + miniGs | 78 | 0.00 | 4.30 | 0.10 | 0.69 | 0.00 | 82.95 | 0.26 | 0.80 | 0.00 |
| | | JSR1-β2AR + miniGs | 88 | 0.00 | 3.60 | 0.03 | 0.68 | 0.00 | 84.23 | 0.16 | 0.81 | 0.01 |
| | | JSR1-DRD1 + miniGs | 111 | 0.00 | 3.72 | 0.02 | 0.72 | 0.00 | 83.51 | 0.14 | 0.82 | 0.00 |
| | | JSR1-JellyOp + miniGs | 76 | 0.00 | 4.45 | 0.02 | 0.57 | 0.00 | 83.99 | 0.21 | 0.74 | 0.01 |
| | | JSR1-LSHR + miniGs | 67 | 0.00 | 5.10 | 0.13 | 0.62 | 0.00 | 76.13 | 0.40 | 0.77 | 0.00 |
| | | JSR1-TAAR9 + miniGs | 92 | 0.00 | 4.72 | 0.03 | 0.66 | 0.00 | 79.90 | 0.10 | 0.80 | 0.00 |

**Table S4. Interface confidence parameters of the AF predictions for the JSR1-WT, JSR1-S199F, different JSR1-S199F-DRD1-chimeras, and DRD1.** The table displays the number of contact points predicted to have a contact probability of > 0.2 and their average (Avg.) predicted aligned error (PAE) for the receptor. Additionally, the average contact probability and the chain-interface predicted template modelling values (ipTM) are shown. The data are colour coded from green to red based on their values. For PAE, lower values indicate better predictions and are shown in green (overall range: 3.01–9.7). In contrast, for number of contacts, avg. contact probability, pLDDT and iPTM, higher values represent better predictions and are displayed in green. The values are given as mean and SD of the five models provided by a single AlphaFold prediction with five seeds.

| Constructs | number of contacts >0.2 | | Avg. Receptor PAE | | Avg. contact probability | | Avg. Receptor pLDDT | | ipTM | |
|---|---|---|---|---|---|---|---|---|---|---|
| | M | SD | M | SD | M | SD | M | SD | M | SD |
| JSR1-WT<br>+ miniGiq04 | 85 | 0.00 | 4.30 | 0.07 | 0.67 | 0.00 | 86.13 | 0.27 | 0.74 | 0.01 |
| JSR1-WT<br>+ miniGs | 97 | 0.00 | 9.12 | 0.53 | 0.43 | 0.00 | 71.78 | 1.04 | 0.56 | 0.00 |
| JSR1-S199F<br>+ miniGs | 79 | 0.00 | 9.73 | 0.18 | 0.40 | 0.00 | 70.25 | 0.43 | 0.53 | 0.02 |
| C1 - JSR1-S199F-DRD1-CT<br>+ miniGs | 76 | 0.00 | 9.50 | 0.15 | 0.41 | 0.00 | 69.96 | 0.35 | 0.54 | 0.01 |
| C2 – JSR1-S199F-DRD1-ICL3<br>+ miniGs | 116 | 0.00 | 4.79 | 0.08 | 0.61 | 0.00 | 79.89 | 0.34 | 0.76 | 0.01 |
| C3 - JSR1-S199F-DRD1-ICL3-CT<br>+ miniGs | 111 | 0.00 | 4.63 | 0.08 | 0.67 | 0.00 | 79.59 | 0.52 | 0.79 | 0.01 |
| C4 - JSR1-S199F-DRD1-ICL3-H8-CT<br>+ miniGs | 110 | 0.00 | 4.30 | 0.07 | 0.69 | 0.00 | 82.43 | 0.32 | 0.79 | 0.01 |
| C5 - JSR1-S199F-DRD1-ICL2-ICL3-H8-CT<br>+ miniGs | 109 | 0.00 | 3.61 | 0.06 | 0.72 | 0.00 | 84.05 | 0.30 | 0.82 | 0.00 |
| DRD1<br>+ miniGs | 100 | 0.00 | 3.39 | 0.03 | 0.75 | 0.00 | 83.79 | 0.18 | 0.84 | 0.00 |

**Table S5.** The table reports the mean and SEM of the parameters extracted from time-resolved curves from different biological replicates. Max_raw normalized was only applied to datasets which were measured in parallel with c5, since c5 was used for normalization. The rate constants ($k_1$- $k_3$) reported here describe the apparent kinetics of the GloSensor cAMP response in HEK293T cells shown in Fig. 3e and three additional biological replicates. The curves were fitted using Scipy. $k_1$ is the rate constant extracted from the rising phase, $k_2$ from the falling phase of cAMP and $k_3$ for the second rising phase when receptors were activated, deactivated and activated again. The chimera c1 had Gi activity (opposite effect to Gs and the rate constants were therefore not fitted. When receptors were activated only once (1x), $k_2$ represents the rate of spontaneous signal decay (grey background) and no $k_3$ is reported. The labels c5-0.5 and c5-2 denote conditions where half or double the standard amount of DNA was used for transfection. All fits and goodness-of-fit parameters are available in optoDRD1_fitting_summary.xlsx, and the fitted curves are displayed in optoDRD1_fitting_plots.pdf. Representative raw data used for this analysis are shown in Figs. 3e, 5a-c, and 6c, and the analyzed data from replicates are plotted in Figs. 3b and S9.

| | basal activity | | | max_raw | | | fold change | | | 30 min after deactivation | | | max_2 | | |
|---|---|---|---|---|---|---|---|---|---|---|---|---|---|---|---|
| **name** | **mean** | **SEM** | **n** | **mean** | **SEM** | **n** | **mean** | **SEM** | **n** | **mean** | **SEM** | **n** | **mean** | **SEM** | **n** |
| **c1** | 1594 | 520 | 4 | 697 | 230 | 4 | 0 | 0 | 4 | 166 | 24 | 4 | 110 | 18 | 4 |
| **c2** | 994 | 172 | 4 | 3859 | 1534 | 4 | 4 | 1 | 4 | 22 | 4 | 4 | 86 | 14 | 4 |
| **c3** | 1030 | 244 | 4 | 34717 | 18657 | 4 | 28 | 11 | 4 | 10 | 2 | 4 | 89 | 13 | 4 |
| **c4** | 1387 | 470 | 4 | 59561 | 31898 | 4 | 35 | 9 | 4 | 22 | 4 | 4 | 80 | 6 | 4 |
| **c5** | 4936 | 1679 | 5 | 183776 | 71473 | 5 | 34 | 8 | 5 | 32 | 8 | 5 | 87 | 8 | 5 |
| **DRD1** | 31261 | 12714 | 5 | 212131 | 55077 | 5 | 43 | 25 | 5 | 11 | | 1 | 97 | 9 | 3 |
| **JellyOp-1** | 4716 | 2386 | 3 | 541529 | 126358 | 3 | 171 | 61 | 3 | 3 | 1 | 3 | | | 0 |
| **c5-0.5** | 3644 | | 1 | 124824 | | 1 | 34 | | 1 | 16 | | 1 | 81 | | 1 |
| **c5-2** | 8052 | | 1 | 208050 | | 1 | 26 | | 1 | 21 | | 1 | 90 | | 1 |
| **n2a-c5-1x** | 4607 | 2451 | 3 | 25776 | 15346 | 3 | 6 | 1 | 3 | | | 0 | | | 0 |
| **n2a-c5** | 4965 | 2402 | 3 | 25030 | 13249 | 3 | 5 | 1 | 3 | 16 | 3 | 3 | 55 | 3 | 3 |

| | max_raw normalized | | | k1 | | | k2 | | | k3 | | |
|---|---|---|---|---|---|---|---|---|---|---|---|---|
| **name** | **mean** | **SEM** | **n** | **mean** | **SEM** | **n** | **mean** | **SEM** | **n** | **mean** | **SEM** | **n** |
| **c1** | 0.006 | 0.001 | 4 | | | 0 | | | 0 | | | 0 |
| **c2** | 0.031 | 0.005 | 4 | 0.404 | 0.044 | 4 | 0.168 | 0.040 | 4 | 0.420 | 0.030 | 4 |
| **c3** | 0.213 | 0.008 | 4 | 0.117 | 0.012 | 4 | 0.098 | 0.015 | 4 | 0.136 | 0.009 | 4 |
| **c4** | 0.353 | 0.013 | 4 | 0.111 | 0.013 | 4 | 0.054 | 0.016 | 4 | 0.117 | 0.014 | 4 |
| **c5** | 1.000 | 0.000 | 5 | 0.142 | 0.020 | 5 | 0.050 | 0.016 | 5 | 0.142 | 0.020 | 5 |
| **DRD1** | 1.060 | 0.154 | 3 | 0.131 | 0.019 | 4 | 0.107 | 0.033 | 4 | 0.124 | 0.005 | 4 |
| **JellyOp-1** | 1.259 | 0.037 | 3 | 0.357 | 0.046 | 3 | 0.090 | 0.008 | 3 | | | 0 |
| **c5-0.5** | 0.803 | | 1 | 0.133 | | 1 | 0.053 | | 1 | 0.120 | | 1 |
| **c5-2** | 1.338 | | 1 | 0.122 | | 1 | 0.057 | | 1 | 0.123 | | 1 |
| **c5-1** | 1.074 | 0.074 | 3 | 0.201 | 0.003 | 3 | 0.003 | 0.002 | 3 | | | 0 |
| **DRD1-1** | | | 0 | 0.103 | 0.008 | 3 | 0.001 | 0.001 | 3 | | | 0 |
| **n2a-c5-1** | | | 0 | 0.248 | 0.044 | 3 | 0.011 | 0.001 | 3 | | | 0 |
| **n2a-c5** | | | 0 | 0.280 | 0.029 | 3 | 0.126 | 0.032 | 3 | 0.276 | 0.020 | 3 |

**Table S6.** The table lists G protein positions (identified via GPCRdb) that are in contact with DRD1 in at least 50% of the available structures (listed in the PDB IDs column, n = 26). Residues within the last 13 amino acids of the Gα C-terminus (α5 helix), which were exchanged in the Gsx assay, are marked in red.

| | G protein position forming receptor interaction in at least 50% of PDBs |
|---|---|
| 1 | G.S1.02 |
| 2 | G.s2s3.02 |
| 3 | G.S3.01 |
| 4 | G.S3.03 |
| 5 | G.H4.13 |
| 6 | G.H4.16 |
| 7 | G.h4s6.03 |
| 8 | G.h4s6.20 |
| 9 | G.S6.03 |
| 10 | G.H5.08 |
| 11 | G.H5.11 |
| 12 | G.H5.12 |
| 13 | G.H5.13 |
| 14 | G.H5.15 |
| 15 | G.H5.16 |
| 16 | G.H5.17 |
| 17 | G.H5.19 |
| 18 | G.H5.20 |
| 19 | G.H5.23 |
| 20 | G.H5.24 |
| 21 | G.H5.25 |
| 22 | G.H5.26 |

| PDB IDs |
|---|
| 7X2D |
| 7CKZ |
| 7LJC |
| 7LJD |
| 7JVQ |
| 7JOZ |
| 7JVP |
| 9I52 |
| 7JV5 |
| 7F24 |
| 7F23 |
| 9I54 |
| 7F1Z |
| 7F1O |
| 7F0T |
| 7CRH |
| 7CKY |
| 7CKX |
| 7X2C |
| 7CKW |
| 9LWC |
| 9LLJ |
| 8IRR |
| 9LLE |
| 9LLF |
| 9LLH |

**References only in supplementary**

1. Stojanovic, T. *et al.* Validation of dopamine receptor DRD1 and DRD2 antibodies using receptor deficient mice. *Amino Acids* **49**, 1101–1109 (2017).
